# Identification of CRYAB^+^ KCNN3^+^ SOX9^+^ astro-like and EGFR^+^ PDGFRA^+^ OLIG1^+^ oligo-like tumoral cells in diffuse low-grade gliomas and implication of Notch1 signalling in their genesis

**DOI:** 10.1101/2021.02.09.430313

**Authors:** Meera Augustus, Donovan Pineau, Franck Aimond, Safa Azar, David Lecca, Nicolas Leventoux, Frédérique Scamps, Sophie Muxel, Amélie Darlix, William Ritchie, Catherine Gozé, Valérie Rigau, Hugues Duffau, Jean-Philippe Hugnot

## Abstract

IDH1-mutated gliomas are slow growing brain tumours, which progress into high-grade gliomas. They present intra-tumoural cell heterogeneity, but no good markers are available to distinguish the different cell subtypes. The molecular mechanisms underlying the formation of this cell diversity is also ill defined. Here we report that the SOX9 and OLIG1 transcription factors, which specifically label astrocytes and oligodendrocytes in the normal brain, reveal the presence of two largely non-overlapping tumoural populations in IDH1-mutated oligodendrogliomas and astrocytomas. Astro-like SOX9^+^ cells additionally stain for APOE, CRYAB, ID4, KCNN3, while oligo-like OLIG1^+^ cells stain for ASCL1, EGFR, IDH1, PDGFRA, PTPRZ1, SOX4, and SOX8. GPR17, an oligodendrocytic marker, was expressed by both cells. These two sub-populations appear to have distinct BMP, NOTCH1, and MAPK active pathways as stainings for BMP4, HEY1, HEY2, p-SMAD1/5 and p-ERK were higher in SOX9^+^ cells. We used primary cultures and a new cell line to explore the influence of NOTCH1 activation and BMP treatment on low-grade glioma cell phenotype. This revealed that NOTCH1 globally reduced oligodendrocytic markers and IDH1 expression while upregulating *APOE, CRYAB, HEY1/2* and an electrophysiologically Ca^2+^-activated apamin-sensitive K^+^ channel (KCNN3/SK3). This was accompanied by reduction in proliferation. Similar effects of NOTCH1 activation were observed in non-tumoural human oligodendrocytic cells, which additionally induced strong SOX9 expression. BMP treatment reduced OLIG1/2 expression and strongly upregulated CRYAB and NOGGIN, a negative regulator of BMP. The presence of astro-like SOX9^+^ and oligo-like OLIG1^+^ cells in diffuse low-grade gliomas raise new questions about their role in the pathology.

## 1. Introduction

Glioma represents nearly 26% of all central nervous system tumours and 80% of primary malignant brain tumours [1]. Based on the histopathological and clinical evaluations, adult diffuse gliomas have been classified into low grade (grade II), anaplastic (grade III) and glioblastomas (GBM, grade IV) [2]. Diffuse low-grade gliomas (DLGG) account for approximately 15% of all gliomas and their incidence rate is about 1/100,000 person-years. These are rare and slow-growing tumours, primarily occurring in young adults [3,4]. Due to their invasive nature, DLGG inexorably relapse 5-10 years after resection, which is often associated with an increase in malignancy leading to rapid patient death [5,6].

Most of DLGG carry a recurrent missense mutation R132H in the gene encoding cytosolic isocitrate dehydrogenase 1 (IDH1) [7]. Mutated cells accumulate the oncometabolite 2-HG leading to cellular differentiation and cell death defect [8,9]. IDH1-mutated DLGG are further sub-grouped into oligodendrogliomas or astrocytomas by the presence or absence of an unbalanced translocation of chromosomes 1 and 19 resulting in the deletion of 1p and 19q (1p/19q co-deletion) [10]. Other genetic alterations often found concomitantly in oligodendrogliomas are CIC, FUBP1, and TERT promoter mutations while astrocytomas most often carry ATRX and p53 mutations [11–13].

In addition to genetic abnormalities, there have been reports suggesting the importance of altered signalling pathways in gliomas. This is well documented in high grade gliomas with the implication of dysregulated receptor tyrosine kinase signallings (epithelial growth factor (EGFR) [14], platelet-derived growth factor (PDGFR) [15] as well as PI3K-AKT-mTOR and Ras-MAPK pathways [16]. However, there are only a handful of studies exploring the signalling aspects in DLGGs [17–20].

It is now well established that most cancers comprise of different tumoural cell subtypes differing for instance in their differentiation, proliferating and metabolic states. DLGG are no exception. Indeed, using immunofluorescence on glioma sections, we previously reported on intratumoural heterogeneity in DLGG [20]. This heterogeneity is also supported by two recent single cell RNA seq studies [21,22]. Additionally, we reported that approximately 20% of DLGG have transformation foci with cells showing a high level of activated STAT3 pathway and reduced level of a lipid metabolism enzyme, ETNPPL [23]. One consequence of this cellular heterogeneity is that current treatments may only target sub-populations of cells while leaving other cell types unaffected and prone to tumor relapse. It is also not known whether all tumoural cell subtypes have the same ability to proliferate and to invade the brain, a major obstacle to treat these tumours. Finally, the intimate molecular mechanisms and pathways underlying the formation of this cellular heterogeneity in DLGG remain ill defined.

Clearly, a better description of this cellular heterogeneity and its formation would certainly help define innovative therapeutic strategies. In particular, simple and reliable markers to reveal this cellular heterogeneity in patients would be very useful in routine pathology practices. In this article, we addressed these pending issues by using freshly resected IDH1-mutated DLGG samples. Immunofluorescence, primary DLGG cell cultures, QPCR and electrophysiology were used to study cellular heterogeneity and active pathways in DLGG. This revealed the presence of two largely non-overlapping astro-like and oligo-like cell populations showing different transcriptional factors, active pathways and receptors. In addition, by conducting functional analysis *in vitro*, we identified Notch1 and BMP pathways as important regulators of DLGG cell phenotype and electrophysiological properties. The presence of two distinct cellular populations in DLGG raises new questions about their role in the pathology, malignant progression and response to treatments.

## 2. Materials and Methods

### 2.1. Patient Samples

Patients used in the article are listed in Table S1. Grade II diffuse low grade gliomas were classified based on WHO criteria (Louis et al., 2016) by a neuropathologist (Pr. V Rigau, Montpellier hospital). Tumours were categorized as 1/ astrocytomas based on IDH1 R132H, p53 stainings and loss of nuclear staining for ATRX by IHC and 2/ oligodendrogliomas based on IDH1 R132H staining and 1p19q codeletion. IDH1 mutation was also confirmed by sequencing of exon 4. 1p/19q co-deletion was assessed by molecular detection of loss of heterozygosity using polymorphic markers within 1p and 19q chromosome arms as described previously [24].

### 2.2. Immunofluorescence on DLGG tissues and cells

Freshly resected tumours were fixed using 4% paraformaldehyde for 1 hour at room temperature (RT) followed by cryopreservation using successive sucrose solutions (10, 20, 30%). Tumours were cryosectioned (10 μm) and sections were washed with 0.1 M PBS-Glycine to reduce background signal. Permeabilization and blocking were performed with 0.3% Triton X-100 and 10% donkey serum (Sigma) followed by overnight incubation with primary antibodies (see supporting information Table No. 4 for the list of antibodies and dilutions) and one hour incubation with Alexa 488- or Cy3-conjugated secondary antibodies (Jackson ImmunoResearch). No primary antibody or antibody against GFP was used as a negative control. Nuclei were counterstained using Hoechst 33342. A minimum of 300 cells was counted to distinguish the two non-overlapping SOX9^+^ and OLIG1^+^ sub-populations (Fig 1A and Fig S2A). A minimum of 100 cells was counted to determine the co-expression of specific markers in SOX9^+^ and OLIG1^+^ cells. For cell cultures, coverslips were fixed using 4% paraformaldehyde for 20 minutes and immunofluorescences were performed as for tumor sections with 0.1% Triton X-100. For staining with O4 antibody, the permeabilization step was omitted. Images were taken using a Zeiss Apotome Z2 microscope.

**Figure 1:**
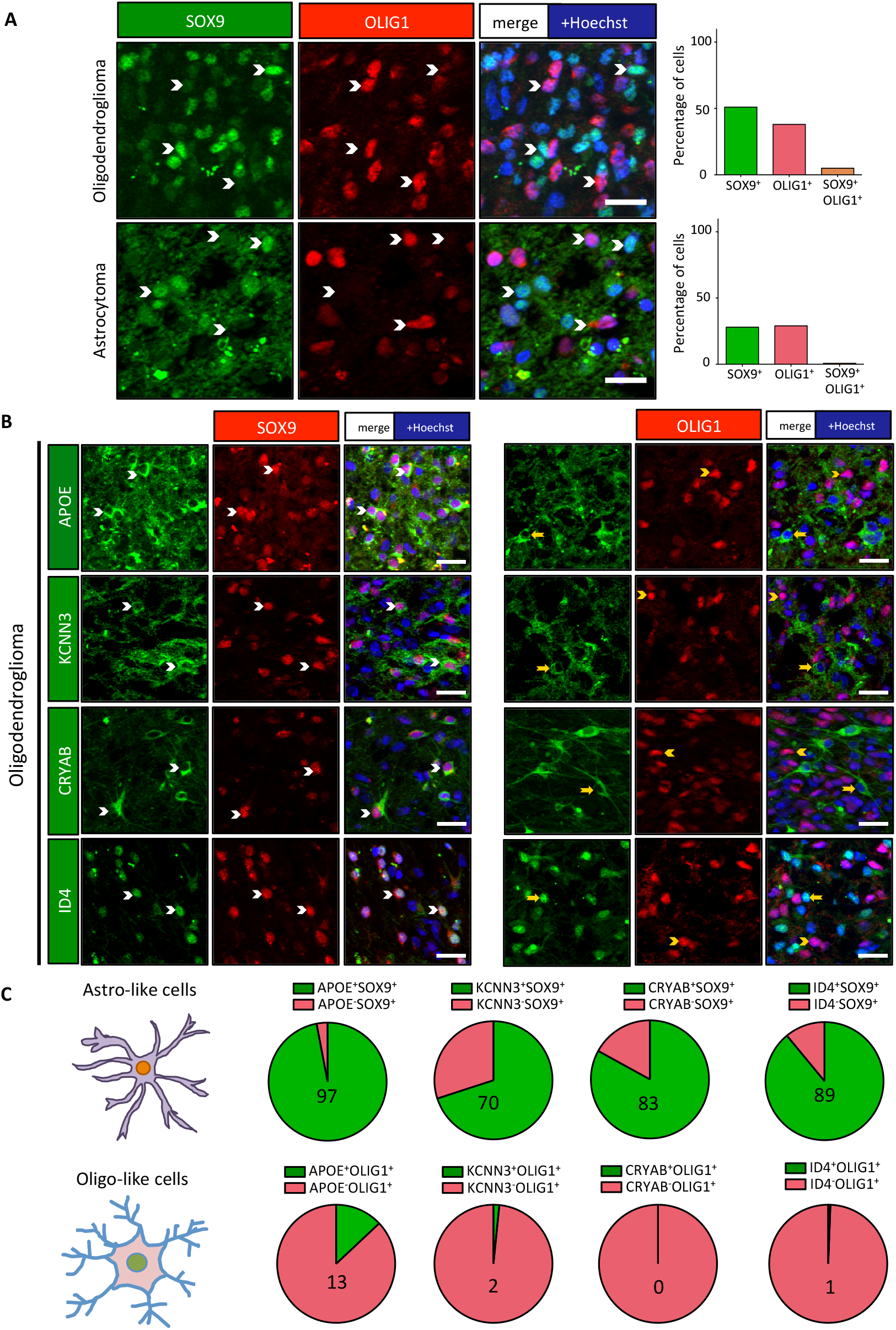
Two non-overlapping cell subpopulations detected in DLGG. **(A)** Immunofluorescence performed on one oligodendroglioma and one astrocytoma using antibodies against SOX9 (green) and OLIG1 (red) revealed the presence of two non-overlapping cell subpopulations. White arrowheads mark cells expressing either SOX9 or OLIG1 alone. Scale bars=20 μm. Bar diagrams show the percentage of SOX9^+^ cells (green), OLIG1^+^ cells (red) and percentage of cells double positive for SOX9^+^ and OLIG1^+^ (orange) among the total number of cells. The two subpopulations are also detected in other patients (see Fig S2A). **(B)** SOX9^+^ cells show specific protein expression. Double stainings for APOE, CRYAB, KCNN3, and ID4 with SOX9 or OLIG1 showed their preferential expression in SOX9^+^ cells. White arrowheads identify double positive cells while yellow arrow-heads/arrows represent single positive cells. Scale bars=20 μm. **(C)** Pie diagrams representing the percentage of double positive (green) and single positive (red) cells in SOX9^+^ (upper lane) and OLIG1^+^ (lower lane) populations. Numbers indicate the percentage of double positive cells.

### 2.3. Cell cultures derived from DLGG resections

Dissociation of resected tumours was done for 30 min at 37°C using Trypsin (Sigma #T4799, 13 mg/ml), Hyaluronidase (Sigma, #H3884, 7 mg/ml) and DNase I (Roche, #10104159001, 10 mg/ml) then stopped by Trypsin Inhibitor (Sigma, #T9003, 50 mg/ml). Cells were passed through a 70 μm cell strainer (Miltenyi Biotech) and purified by a two-step percoll (Sigma, #GE17-0891-02) density gradient to remove myelin and red blood cells. Microglia cells were removed and O4^+^ cells were collected by sorting with CD11b^+^ and O4^+^ magnetic microbeads respectively (Miltenyi Biotech) according to the manufacturer’s protocol. O4^+^ cells were grown on culture vessels coated with poly-D-lysine (Sigma, #P7886; 25 μg/ml) and laminin (Sigma, #L2020, 2 μg/cm2). Cells were cultured in DMEM/F12 1:1 media (Gibco) supplemented with N2 (Thermo Fisher, #17502048), D-glucose (Sigma, 0.6%), L-glutamine (Thermo Fisher, #25030024, 2mM), B27 w/o vitamin A (Invitrogen), EGF (Peprotech, 20 ng/ml), FGF2 (Peprotech, 10 ng/ml), PDGFA (Peprotech, 20 ng/ml) and heparin (Sigma, 2 μg/ml). The LGG275 cell line was isolated by serial passaging of a DLGG culture showing proliferative cells which was derived from a ATRX/IDH1 R132H mutated resection (obtained from the Montpellier hospital biological resource bank and authorization from the Montpellier hospital Institutional Review Board (IRB ID: 198711) N° IRB-MTP_2019_IRB_MTP_10_15).

### 2.4. Lentivirus infection, RNA extraction and QPCR

O4^+^ purified cells and LGG275 cells were infected with either a control-YFP or NICD-YFP virus (multiplicity of infection=6). NICD cDNA codes for amino acids 1762 to 2556 of human Notch1 [25]. Five days after transduction, RNAs were extracted using Arcturus picopure RNA kit (Thermo Fisher) and 100-200 ng of cDNA was synthesized with random primers and reverse transcriptase (Promega, GoScript). Quantitative PCR was performed using 2.5 ng of cDNA in duplicates. Primers (Sigma) are listed in Supplementary Table No. 3. KAPA SYBR PCR kit (Sigma) was used for QPCR (Light Cycler 480, Roche). Gene expression was calculated using 2-ΔΔCt method using β-actin (ACTB) for normalization. For inhibitor experiments, LGG275 cells were treated with γ-secretase inhibitors (DAPT (GSI-IX, Selleck Chem) and LY411575 (Peprotech)) at 10 mM for 5 days before RNA was extracted for QPCR. For DLL4 ligand experiment (Fig. S13A), plates were coated with DLL4 (5 μg/ml) (Peprotech) or BSA at 4°C overnight before seeding the cells. RNAs were extracted after 4 days. For BMP signalling experiments (Fig. 6C), cells were treated with 10 ng/ml of BMP2 or BMP4 (Peprotech) for 5 days with addition of new cytokine every 2 days.

### 2.5. Electrophysiology

KCNN3 channel currents were measured in LGG275 cells transduced with control-YFP or NICD-YFP viruses, 3 days after the infection. For electrophysiology, cells were placed in bathing (extracellular) solution composed of 147 mM NaCl, 5 mM KCl, 2 mM CaCl2, 1.5 mM MgCl2, 10 mM HEPES, 10 mM glucose, pH 7.4. Recording pipettes filled with a solution containing 135 mM K-methane-sulfonate, 0.1 mM EGTA, 8 mM KCl, 10 mM HEPES, 2 mM MgATP, 0.5 mM NaGTP, pH 7.3 were sealed to the cell membrane. SK3 current was recorded by applying 600 ms electrical ramp from −80 mV holding potential to gradually increasing up to +60 mV in both control-YFP and NICD-YFP transduced cells. Ionomycin-induced and apamin-sensitive SK3 current densities were measured by adding 10 mM ionomycin and 1 mM apamin to the bathing solution respectively. All recordings were performed at RT using an Axopatch 200B amplifier and a Digidata 1322A A/D board (Molecular Devices) and acquired at 5 kHz. Current analysis was performed using Clampfit version 10 (Axon Instruments).

### 2.6. Western blot analysis

Tissue samples were lysed using RIPA lysis buffer (Sigma) consisting of 5 mM NAF, 0.5 mM Na-vanadate, and 13-protease inhibitor cocktail (Roche). Samples were incubated on ice for 30 minutes, centrifuged for 20 minutes at 13,000 rpm at 4°C and the protein concentration in the supernatant was measured with an DC protein assay (BioRad). Samples were separated by SDS-PAGE, transferred on a 0.2 μm PVDF membrane (BioRad). After blocking with 5% non-fat dried milk in TBST (Tris-buffered saline, 0.1% Tween 20), primary antibodies listed in Supplementary Table No. 4 were incubated overnight at 4°C and revealed using peroxidase-conjugated antibodies and an ECL kit (BioRad). Images were captured using ChemiDocTM XRS Imaging system (Biorad).

### 2.7. Statistical Analysis

All experiments were performed at least three times for confirmation. Data are represented as mean +/− standard error of mean (S.E.M.). Statistical differences in experiments were analyzed with tests indicated in the legends using GraphPad 6 Prism software. Unpaired two-tailed t-tests with equal standard deviation were used for QPCR analysis. *, **, *** represent p < .05, <0.01, and <0.001 significance respectively.

## 3. Results

### 3.1. Two distinct populations of astrocytic-like and oligodendrocytic-like tumoural cells in DLGG

In order to distinguish several cell populations in DLGG tumours, we examined three astrocytoma tumours with IDH1 R132H and ATRX mutations and three oligodendroglioma tumours with IDH1 R132H and 1p19q codeletion (Table S1). To identify two potential cell types in these samples, we used OLIG1 and SOX9 as well-known markers for oligodendrocytic and astrocytic-lineage cells respectively (Fig. S1A, B). Fig. 1A and Fig. S2A shows that in the 6 examined tumours, two largely non-overlapping populations of OLIG1^+^ and SOX9^+^ cells were present. To ascertain that these 2 cell populations were indeed tumoural, we performed co-stainings for SOX9 and OLIG1 with (i) ATRX for one astrocytoma and (ii) a specific antibody for the mutated form of IDH1 (IDH1 R132H) in one oligodendroglioma. Fig. S2B shows that in the studied astrocytoma, no nuclear staining for ATRX was observed in the majority (>90%) of OLIG1^+^ and SOX9^+^ cells which are thus tumoural (Fig. S2B, C). For the oligodendroglioma, >90% of OLIG1^+^ cells were co-stained with IDH1 R132H whereas only around 15% of SOX9^+^ cells showed a faint IDH1 R132H staining (Fig. S2B, C). However, as indicated below, this number is probably an underestimation, as the level of expression of the IDH1 enzyme appears lower in SOX9^+^ than in OLIG1^+^ cells.

To ascertain that SOX9^+^ cells show some astrocytic features, further stainings were performed. We selected proteins that are expressed in DLGG tumours and for which literature and database analyses (Fig. S1A, B and Table S2) show a clear preferential expression in the astrocytic lineage. We chose APOE, ID4, KCNN3 astrocytic proteins for which reliable antibodies were used in combination with OLIG1 and SOX9 transcription factors. We also analyzed the expression of CRYAB (alpha-B crystallin) as this protein was described to be elevated up to 22-fold in IDH1-mutated tumours [26]. Stainings for these 4 proteins were performed in one oligodendroglioma and one astrocytoma mutated for IDH1 along with 1p/19q co-deletion or ATRX mutation respectively. Results presented in Fig. 1B, C (oligodendroglioma) and Fig. S3 (astrocytoma) showed that the expression of these 4 proteins were more associated with SOX9^+^ cells than OLIG1^+^ cells.

We then performed the same analysis with proteins that are hallmarks of the oligodendrocytic lineage, namely GPR17, PDGFRA and SOX8 (Fig. S1A, B and Table S2). PDGFRA and SOX8 was clearly more associated with OLIG1^+^ cells in both oligodendroglioma and astrocytoma (Fig. 2A, B and Fig. S4A, B). For GPR17, its expression was more confined to OLIG1^+^ cells in the oligodendroglioma (Fig. 2A, B) whereas a similar expression was found in the SOX9^+^ and OLIG1^+^ subpopulations in the explored astrocytoma (Fig. S4A, B).

**Figure 2:**
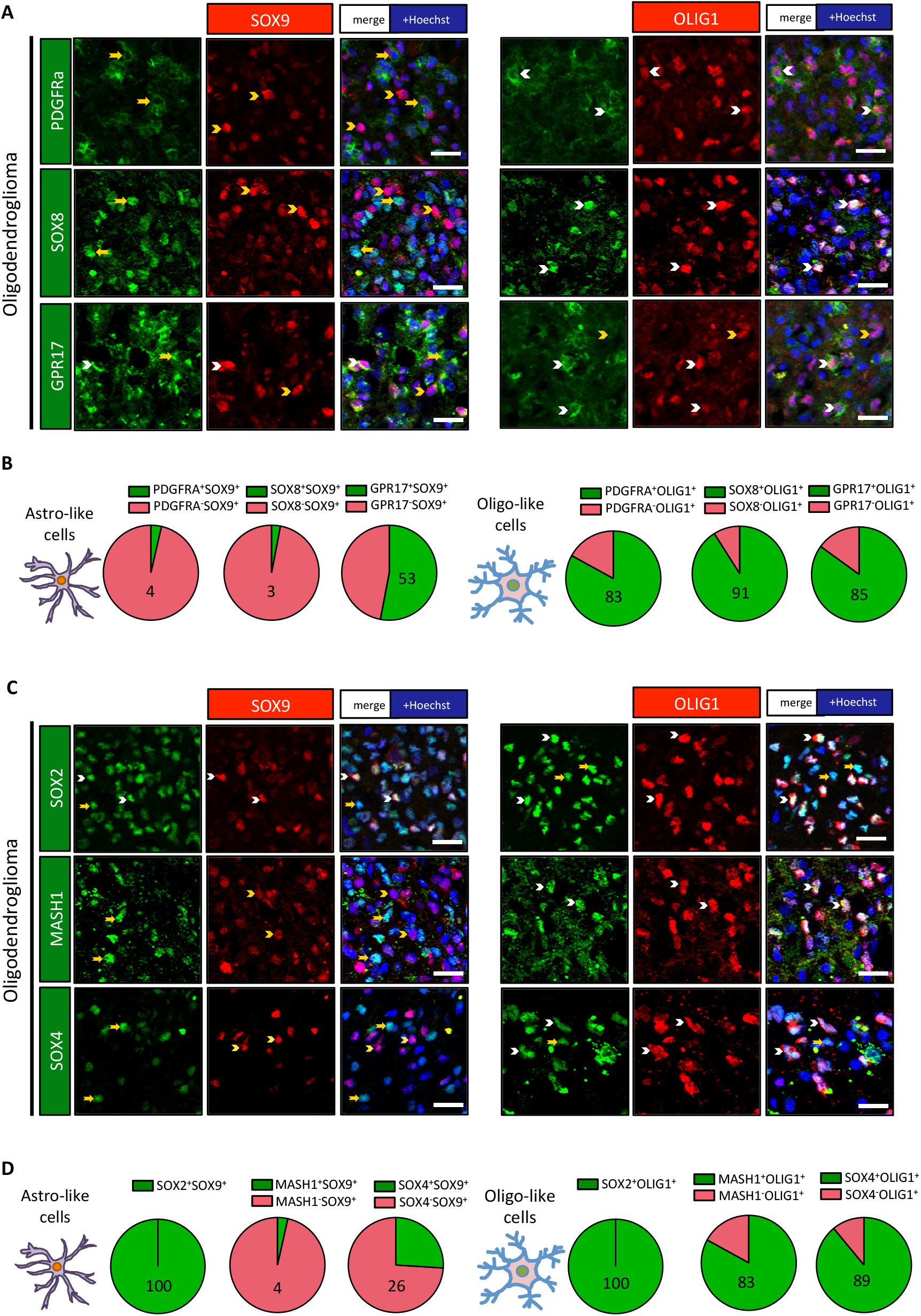
OLIG1^+^ cells express proteins associated to the oligodendrocyte lineage and neural precursor cells. Double immunofluorescences for indicated proteins on one oligodendroglioma. White arrowheads identify double positive cells while yellow arrowheads/arrows represent single positive cells. Scale bars=20 μm. (A) Double stainings for GPR17, PDGFRα, SOX8 with SOX9 or OLIG1 revealed their preferential association with OLIG1^+^ cells in one oligodendroglioma. (B, D) Pie diagrams representing the percentage of double positive (green) and single positive (red) cells in SOX9^+^ and OLIG1^+^ populations. Numbers indicate the percentage of double positive cells. (C) Double stainings for MASH1, SOX2, and SOX4 with SOX9 or OLIG1 revealed the preferential association of MASH1 and SOX4 with OLIG1^+^ cells in one oligodendroglioma while SOX2 is expressed by both populations.

Collectively, these results indicate the presence of at least two cell populations in DLGG showing an astro- and oligo-like phenotype.

### 3.2. Astro- and oligo-like cells have different levels of receptors and signalling proteins

Considering the distinct phenotype of SOX9^+^ and OLIG1^+^ cells, we reasoned that these cells might present different active pathways, receptors and neural developmental transcription factors. We started addressing this issue by performing co-stainings for SOX9 and OLIG1 with other transcription factors which are highly expressed by neural precursor cells during brain development (ASCL1/MASH1, SOX2, SOX4). We found that SOX2 was expressed at the same level in both cell types (Fig. 2C, D and Fig. S4C, D). On the contrary, MASH1 and SOX4 expression was clearly more expressed in OLIG1^+^ cells both in the explored oligodendroglioma (Fig. 2C, D) and astrocytoma (Fig. S4C, D) samples.

We next studied the expression of proteins that are considered as good readouts of activation of BMP, Notch1, and ERK/MAP pathways. We previously reported that in glioblastomas, Notch1 pathway activation led to the upregulation of HEY1 and HEY2 transcription factors [27]. This observation led us to assess HEY1 and HEY2 stainings in the two cellular populations. Results presented on Fig. 3A, D for one oligodendroglioma and Fig. S5A, D for one astrocytoma, indicate that HEY1 and HEY2 were preferentially expressed in SOX9^+^ cells suggesting that Notch1 signalling could be active in these cells. With regards to the BMP signalling, this pathway is often associated with the generation of astrocytes during brain development and in several pathological situations [28,29]. We looked for activation of this pathway by performing co-stainings for SOX9 and OLIG1 with the phosphorylated form of SMAD1/5 (p-SMAD1/5), a signalling protein downstream of BMP pathway. We found that p-SMAD1/5 was preferentially detected in the SOX9^+^ population (Fig. 3B, D and Fig. S5B, D). This suggested that DLGG might express BMP RNA and protein. Indeed, mining of two databases indicates a significant upregulation of BMP2 in grade II/III oligodendrogliomas and astrocytomas compared to non-tumoural tissues and BMP4 was overexpressed in oligodendrogliomas in one database (Fig. S6A). Correlation analysis also showed that BMP2 and BMP4 expression are positively correlated in these tumours (Fig. S6B). We confirmed this result in our samples by QPCR and WB for BMP2 and 4. Results presented in Fig. S7A, B show significant upregulation of both BMP2 and BMP4 mRNA in the 5 explored samples and their strong correlated expression. BMP2 and BMP4 protein were detected by western blot analysis in all examined grade II/III tumours (Fig. S7C). To study which cell types express BMP2 and 4 in DLGG, we performed IF for these proteins. We could not obtain a reliable staining for BMP2 using different antibodies, however we detected a preferential expression of BMP4 in astrocytic SOX9^+^ cells (Fig. 3B, D for one oligodendroglioma and Fig. S5B, D for one astrocytoma).

**Figure 3:**
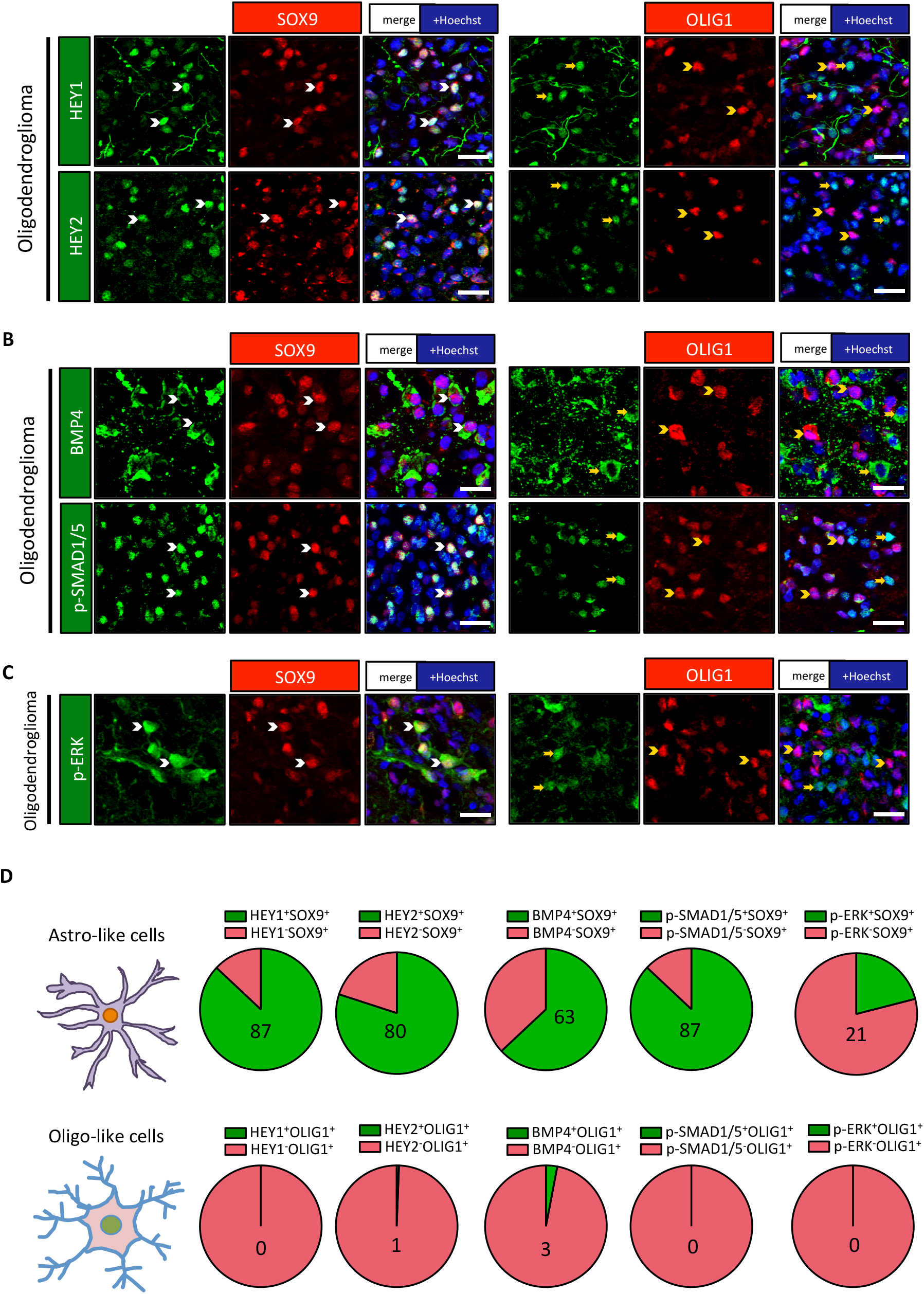
SOX9^+^ cells express specific signalling proteins and transcription factors. (A, B, C) Double immunofluorescences in one oligodendroglioma for HEY1, HEY2, BMP4, p-SMAD1/5 and p-ERK with SOX9 and OLIG1 revealed their preferential expression in SOX9^+^ cells. White arrowheads identify double positive cells while yellow arrowheads/arrows show single positive cells. Scale bars=20 μm. (D) Pie diagrams representing the percentage of double positive (green) and single positive (red) cells in SOX9^+^ and OLIG1^+^ cells. Numbers indicate the percentage of double positive cells.

The receptors for EGF (EGFR) and PTN (PTPRZ1) have important roles in the genesis of gliomas [19,30–35]. We thus examined their expression in the two cell populations. Surprisingly, we found that these two receptors were more expressed in OLIG1^+^ cells in the explored oligodendroglioma (Fig. 4A, B) and astrocytoma (Fig. S8A, B). Activation of EGFR and other receptors activate the ERK/MAPK pathway leading to phosphorylation of the ERK protein (p-ERK). We thus questioned the presence of p-ERK in the two populations by IF. Results presented on Fig. 3C, D and Fig. S5C, D shows that the SOX9^+^ population appears to have higher activation of ERK/MAPK pathway.

**Figure 4:**
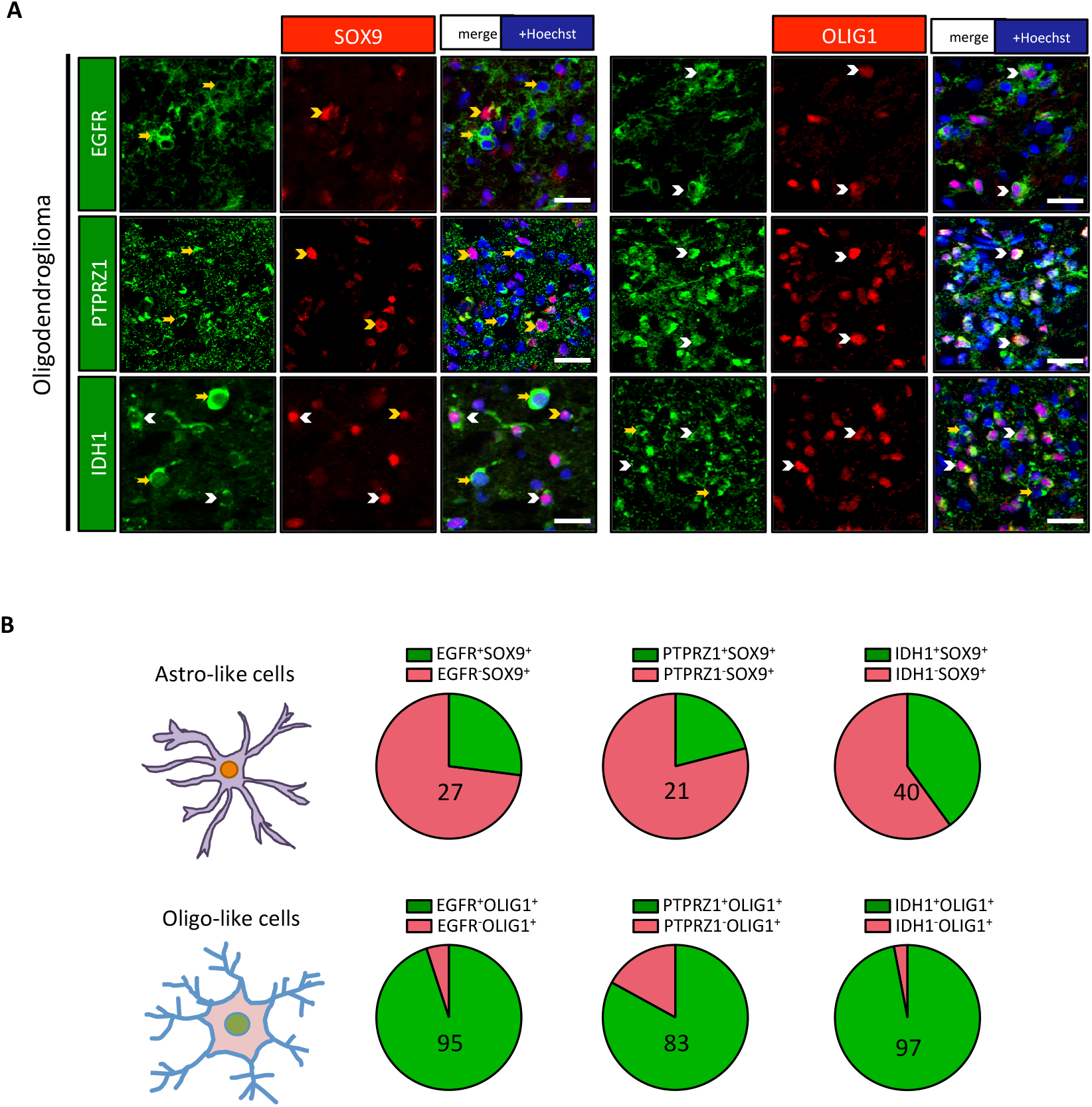
OLIG1^+^ cells express specific receptors and IDH1. (A) Double immunofluorescences in one oligodendroglioma for EGFR, PTPRZ1 and IDH1 with SOX9 and OLIG1 revealed their preferential expression with OLIG1^+^ cells. White arrowheads identify double positive cells while yellow arrowheads/arrows represent single positive cells. Scale bars=20 μm. (B) Pie diagrams representing the percentage of double positive (green) and single positive (red) cells in SOX9^+^ and OLIG1^+^ cells. Numbers indicate the percentage of double positive cells.

All together, these results show that the two identified cell populations in DLGG express different level of receptors and signalling pathways.

### 3.3. Astro- and oligo-like cells express different level of IDH1 enzyme

IDH1 is a cytoplasmic enzyme that converts isocitrate into alpha-ketoglutarate [36]. Unexpectedly, exploration of mouse and human brain single cell databases shows that the IDH1 gene is preferentially expressed by oligodendrocytic lineage cells (Fig. S9). This observation prompted us to explore the presence of IDH1 enzyme in SOX9^+^ and OLIG1^+^ cells by IF with an antibody raised against the wild type enzyme. Fig. 4A, B shows that in the explored oligodendroglioma IDH1 staining was preferentially associated with the OLIG1^+^ cell population. In contrast, in the studied astrocytoma, the expression of IDH1 was similarly detected in both OLIG1^+^ and SOX9^+^ populations (Fig. S8A, B).

### 3.4. Notch1 activation modifies DLGG cells phenotype and reduces their proliferation

The identification of at least 2 clearly distinct cell types in DLGG questioned how these cells are generated. We hypothesized that the Notch1 and BMP pathways may influence DLGG cell phenotype especially as we found that SOX9^+^ and OLIG1^+^ cells have a differential expression of proteins typically involved in these signalling (HEY1/2 and p-SMAD1/5). To evaluate this possibility, we derived primary cultures from DLGG resections mutated for ATRX and IDH1 R132H (Fig. S10). These cultures were established by magnetic sorting for the O4^+^ antigen that is expressed both by immature oligodendrocytes and bipotent astro-oligodendrocyte progenitors in the brain [37]. The presence of tumoural cells in the cultures was evaluated by IF against IDH1 R132H and ATRX. Resected samples can contain both tumoural and non-tumoural territories in variable proportion especially in supratotal resections [38] and consequently, we found that the percentage of IDH1 R132H^+^ ATRX^−^ cells was highly variable between cultures, ranging from 5% to 90%. When few mutated cells were present, the cultures mainly consisted of highly-branched cells (Fig. S11A, B) with small nuclei (perimeter =22.9μm ± 0.3, n=200 cells) expressing high level of ATRX, CNP, OLIG1, O4 and SOX10 whereas stainings for IDH1 R132H, EGFR, GFAP and SOX9 were rare or absent in these cells (Fig. S11C, D). Considering their morphology and their markers, these cells are very likely to be non-tumoural oligodendrocytic cells. In contrast, in cultures containing lots of tumoural cells (Fig. S10A, B), we noted that these have a larger and often abnormal nucleus (perimeter =30.7 μm ± 0.4, n=200 cells), and express high levels of EGFR, OLIG1, SOX9 and weak stainings for CNP and SOX10 (Fig. S10C, D). Compared to the in vivo situation, we could not identify two clearly distinct SOX9^+^ and OLIG1^+^ tumoural cells in these cultures and the majority of cells co-express OLIG1 and SOX9 (Fig. S10C; lane 4). This could be due to phenotypic modifications induced by cell culture conditions as observed for instance in cultured neural stem cells [39].

To test the influence of Notch1 pathway on DLGG cells, we selected 4 independent cultures containing >70% of IDH1 R132H and ATRX mutated cells. Cells were infected with a lentivirus expressing the constitutively active form of Notch1 (Notch intracellular domain, NICD) and YFP, which resulted in strong upregulation of NOTCH1 mRNA expression (Fig. 5A). We analyzed by QPCR the expression of most of the genes that we found differentially expressed in SOX9^+^ and OLIG1^+^ cells on DLGG sections. These are EGFR, IDH1, OLIG1, PDGFRA, PTPRZ1 and SOX4/8 for OLIG1^+^ cells and APOE, CRYAB, HEY1/2, KCNN3, SOX9 for SOX9^+^ cells. OLIG2 gene was also included in the analysis as this transcription factor is often co-expressed with OLIG1 in oligodendrocytic cells [40]. Results presented on Fig. 5A indicate that 7/8 of the genes associated with OLIG1^+^ cells were significantly reduced by NICD expression (IDH1, OLIG1/2, PDGFRA, PTPRZ1, SOX4/8) while 3/6 of the genes preferentially expressed in SOX9^+^ cells (HEY1/2, KCNN3) were significantly upregulated. No significant influence of NICD on SOX9 was detected which was in fact already highly expressed in these cultures (Fig. S10C). APOE showed upregulation (fold change range= 1.3- 2.6) in 3 of 4 explored primary cultures but did not reach significance.

**Figure 5:**
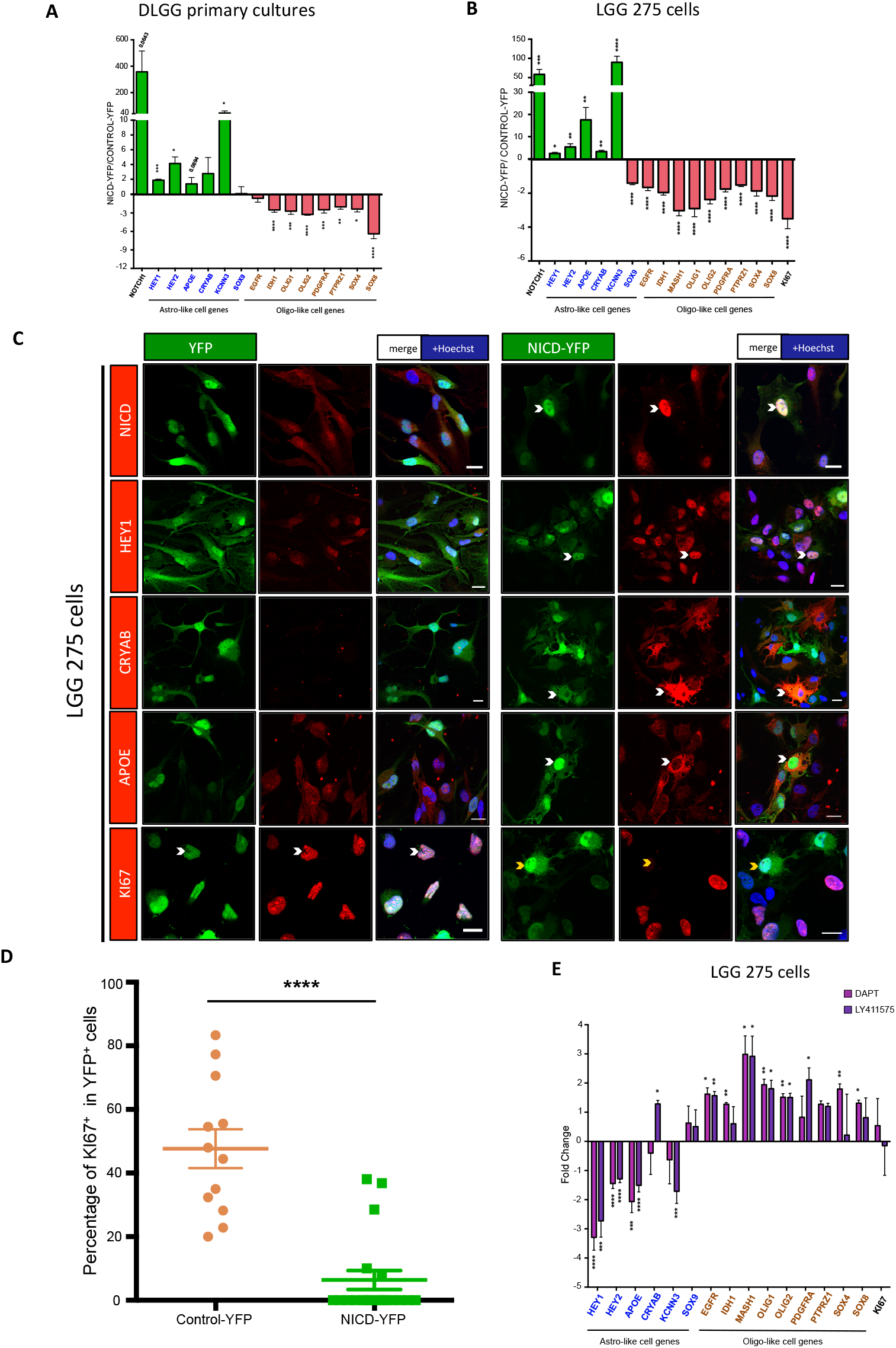
Notch1 activation modifies DLGG cell phenotype and reduces proliferation. (A, B) QPCR analysis for indicated genes in O4-purified primary cultures containing a high % of tumoural cells (>70%) (n=4 patients, astrocytomas) (A) and in LGG275 cells (n=5 independent experiments) (B). Values represent the mean +/−S.E.M. of gene expression fold change observed in cells transduced with NICD-YFP vs. YFP lentiviruses. Genes in blue and brown are markers found preferentially associated respectively with SOX9^+^ and OLIG1^+^ cells on DLGG sections. Tests=two-tailed t-tests. (C) Immunofluorescence for indicated proteins in YFP or NICD-YFP transduced LGG275 cells. White and yellow arrowheads show YFP^+^ cells, which are positive or negative for the assessed protein respectively. Note the nuclear localization of Notch1 activated form (NICD) after transduction with NICD-YFP lentivirus. Scale bars=20 μm. (D) Quantification of MKI67^+^ cells in LGG275 cells transduced with YFP and NICD-YFP lentiviruses. Values represent the mean +/−S.E.M. of percentage of MKI67^+^ cells observed in YFP^+^ cells (n=12 fields, 2 independent experiments). Test=two tailed t-test. (E) QPCR analysis for indicated genes in LGG275 cells treated with Notch1 signalling inhibitors (DAPT, LY411575, 10 μM) for 5 days (n=4 independent experiments). Values represent the mean +/−S.E.M. of gene expression fold change observed in treated vs. control cells. Tests=two tailed t-tests. Genes in blue and brown are markers found preferentially associated respectively with SOX9^+^ and OLIG1^+^ cells on DLGG sections. Tests=two tailed t-tests.

To confirm these results, we used a cell line (named LGG275), which was derived from a DLGG patient with IDH1 and ATRX mutations (Table S1). This cell line contains bipolar and multipolar cells (Fig. S12A) expressing IDH1 R132H but not ATRX (Fig. S12 B lane 1,3) and grow very slowly (doubling time=9.6 +/− 0,1 days, n=6 wells). Fig. S12B, C shows that LGG275 cells express both SOX9 and OLIG1 proteins (lane 4) together with EGFR, CNP and SOX10 (lane 1-3) as seen in primary tumoural cultures (Fig. S10). These cells were transduced with YFP and YFP-NICD lentivirus, which led to strong upregulation of NOTCH1 RNA (Fig. 5B) and a clear nuclear localization of the NICD fragment (Fig. 5C). NICD in these cells downregulated 9/9 of oligodendrocytic genes (ASCL1/MASH1, EGFR, IDH1, OLIG1/2, PDGFRA, PTPRZ1 and SOX4/8) and upregulated 5/6 of astrocytic genes (APOE, CRYAB, HEY1/2, KCNN3). SOX9 was moderately but significantly downregulated by NICD in this cell line. We could confirm the upregulation of APOE, CRYAB, HEY1, and KCNN3 proteins by NICD by IF (Figs. 5C and 6A). In order to explore the effect of Notch1 activation in a more physiological setup, we exposed LGG275 cells to DLL4, a Notch1 ligand predominantly expressed in endothelial cells [41,42]. This resulted in the significant upregulation of APOE, HEY1 and KCNN3 while the expression of all oligodendrocytic genes was reduced (Fig. S13A). The influence of Notch1 signalling was also monitored using two γ-secretase inhibitors (DAPT and LY411575) that blocks Notch1 receptor cleavage thereby preventing signal activation. Fig. 5E shows that these inhibitors downregulate APOE, HEY1/2, and KCNN3 expression while globally increasing oligodendrocytic genes (ASCL1/MASH1, EGFR, IDH1, OLIG1/2, PDGFRA, SOX4/8) (Fig. 5E).

**Figure 6:**
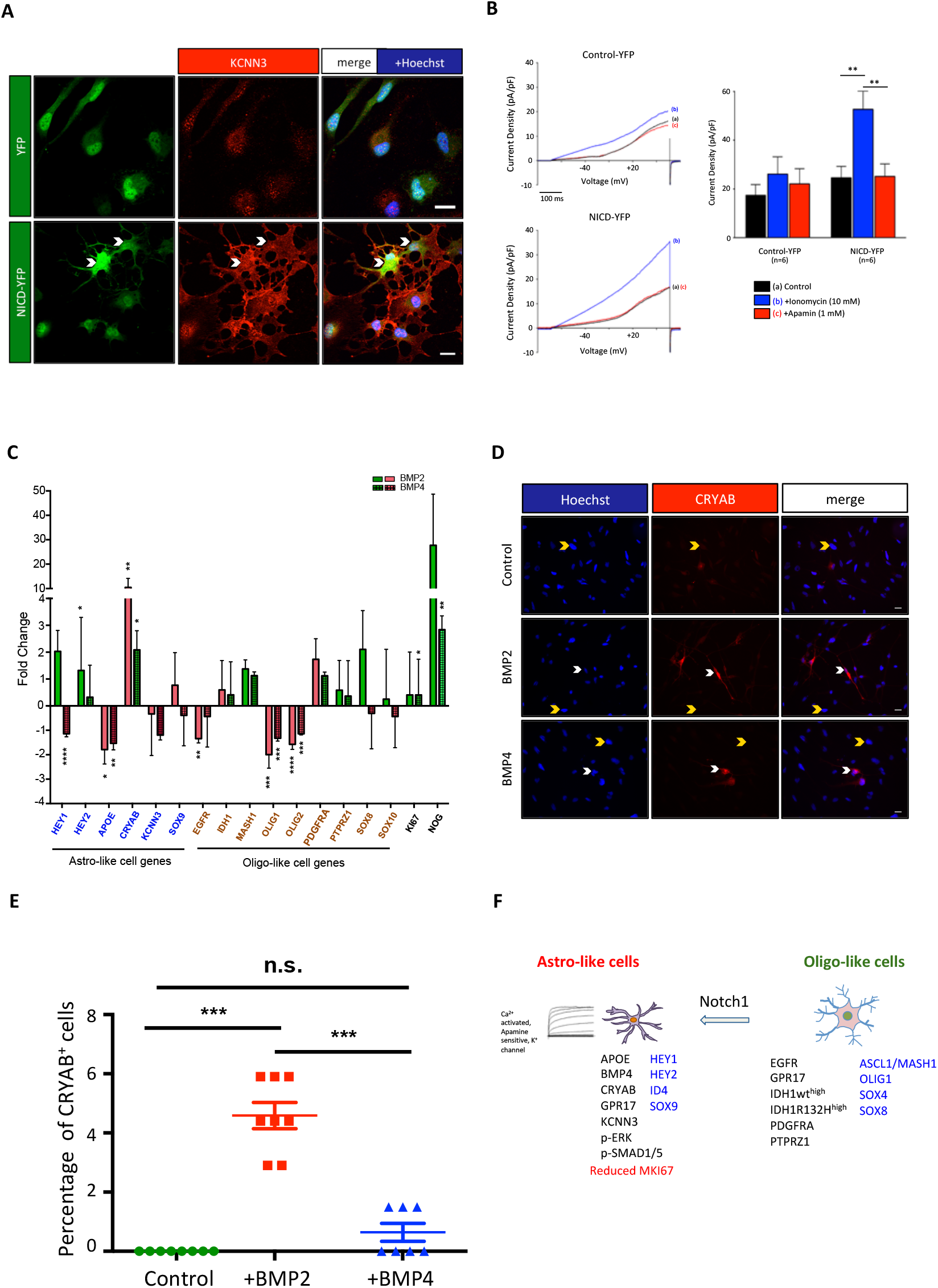
Notch1 induced KCCN3/SK3 channels. BMP influence on DLGG cell phenotype. (A) Immunofluorescence for KCNN3/SK3 channels in YFP and NICD-YFP transduced LGG275 cells. Changes in cellular morphology upon Notch1 activation is also visible. Scale bars=20 μm. (B) KCNN3/SK3 currents induced by Notch1 signalling. Channel activity was studied in YFP^+^ cells using the whole-cell patch clamp technique under voltage clamp configuration. Left panels: Representative currents recorded by applying a 600 ms electrical ramp from −80 mV holding potential to +50 mV in control-YFP (top) and NICD-YFP transduced (bottom) cells in the 3 indicated conditions (a, b, c). An increase in current density was observed in the presence of 10 μM ionomycin (to increase intracellular Ca2^+^ level) in YFP-NICD transduced cells (blue curve). This current increase was specifically blocked by 1 μM apamin (a specific SK channel blocker) (red curve). Right panels: Bar charts showing the means ± S.E.M. of current densities obtained at +40 mV from current/voltage curves obtained after 500 ms voltage steps from −80 mV holding potential to +40 mV every 10 mV in YFP and YFP-NICD transduced cells. Ionomycin-induced and apamin-sensitive current densities were significantly higher in YFP-NICD transduced cells compared to control cells (n=6 cells, 3 independent experiments, test=two-tailed t-test). (C) QPCR analysis for indicated genes in LGG275 cells treated for BMP2 or BMP4 (10 ng/ml) for 5 days (n=4 independent experiments). Values represent the mean +/−S.E.M. of gene expression fold change observed in treated vs. control cells. Genes in blue and brown are markers found preferentially associated respectively with SOX9^+^ and OLIG1^+^ cells on DLGG sections. Tests=two-tailed t-tests. (D) Immunofluorescence for CRYAB in control or BMP2/4 treated LGG275 cells. White and yellow arrowheads show CRYAB^+^ and CRYAB^−^ cells respectively. Scale bars=10 μm. (E) Quantification of CRYAB^+^ cells after BMP2/4 treatment. Values represent the mean +/−S.E.M. of percentage of CRYAB^+^ cells in control or treated cells. (n=7 fields, 3 coverslips). (F) Graphical summary of main results. Transcription factors are in blue.

Finally, as we previously found that Notch1 activation drastically reduced the rate of proliferation of glioblastoma stem cells [27], we also assessed the effect of NICD on the proliferation of LGG275 cells by measuring the expression of the proliferation marker MKI67 by IF and QPCR. As presented on Fig. 5B, C and D, we observed that NICD expression led to a sharp reduction in proliferation.

Collectively, these results show a major role for Notch1 signalling in controlling phenotype and proliferation of DLGG cells.

### 3.5. Notch1 activation modifies electrophysiological properties of DLGG cells

Among the astrocytic genes upregulated by Notch1 activation, the strongest effect was observed for KCNN3 with a fold change of 30 and 100 in primary DLGG cells and the LGG275 line respectively. KCNN3 (also known as SK3) is a small conductance K^+^ channel activated by Ca2^+^, which can be specifically blocked by the bee venom apamin [43]. This channel is expressed in the brain but its role in astrocytes and gliomas is currently unknown. To see whether NICD could elicit KCNN3/SK3 currents, we performed electrophysiology on control-YFP or NICD-transduced LGG275 cells. Results presented on Fig. 6B, showed that in NICD-transduced cells, increase of Ca2^+^ by ionomycin led to an elevated outward K^+^ current which is blocked by apamin. This indicates that NICD can induce a functional expression of KCNN3 channels in DLGG cells.

### 3.6. Notch1 activation induces SOX9 in human oligodendrocytic cells

We were surprised not to observe an upregulation of SOX9 by Notch1 activation in DLGG tumoural primary cultures and in LGG275 cells (Figs. 5A, B) as this gene is upregulated by Notch1 in other contexts [44]. We reasoned that in fact, the high level and maybe already saturated expression of SOX9 in these cells (Fig. S10C and S12B) might preclude an upregulation after NICD transduction. To test this hypothesis, we used 3 primary cultures derived from DLGG supratotal resections, which contained mostly (>90%) non-tumoural SOX10^high^ oligodendrocytic cells (Fig. S11). In these cultures, SOX9 could be barely detected by IF (Fig. S11C; lane 6). QPCR for SOX9 validated this low level of expression compared to tumoural DLGG primary cultures and LGG275 cells (Fig. S13B). Fig. S13E shows that overexpression of NICD in these cells, induced a sharp upregulation of SOX9 (fold change=14.8) that could also be observed at the protein level by IF (Fig. S13C, D). As observed in LGG275 and tumoural DLGG primary cultures, NICD overexpression in these cells also reduced oligodendrocytic genes expression while increasing APOE, CRYAB, HEY1/2 and KCNN3 (Fig. S13E).

### 3.7. BMPs influence on DLGG cell phenotype

BMP proteins are important regulators of neural precursor fate by controlling their phenotype and proliferation [28]. BMP can also induce astrocytic differentiation of oligodendroglioma propagating cells [29]. Their effect not only hinge on the canonical Smad pathway but also by interacting with the Notch1 pathway and HEY1/2 transcription factors [45]. The high expression of BMP proteins found in DLGG tumours especially in astro-like cells (Fig. 3B, D and Fig. S5B, D) prompted us to determine the effect of BMP2/4 on LGG275 cell phenotype. The cells were treated with BMP2 or 4 for 5 days and their phenotype was assessed by QPCR as done for Notch1. Results in Fig. 6C show that the effects were much less contrasted than for Notch1 and only OLIG1/2, and APOE were significantly downregulated by both cytokines. One remarkable variation was observed for CRYAB, which was upregulated 10 and 2 fold by BMP2 and BMP4 treatment respectively (Fig. 6C). Correspondingly, the CRYAB protein, which is not detected in control LGG275 cells by IF, become clearly present in a fraction of cells exposed to BMP2 and to a lesser extent to BMP4 (Fig. 6D, E). Last, mining of the DLGG TCGA glioma databases show that expression of BMP2/4 is highly correlated to the expression of the BMP-inhibitor NOG (NOGGIN) gene (Fig. S6A, B). We confirmed this tight correlation by performing QPCR for NOG in the five DLGG resections we previously analyzed for BMP2/4 (Fig. S7A, B). This suggested that BMP2 or 4 cytokines might control NOG expression in DLGG. Indeed, using QPCR, we found that BMP2 was a powerful regulator of NOG in LGG275 cells with a 30-fold increase (Fig. 6C). BMP4 could also significantly upregulate NOG however only 3-fold (Fig. 6C).

## 4. Discussion

In this article, we studied tumoural cell heterogeneity in IDH1-mutated diffuse low-grade gliomas, a subtype of gliomas for which little is known at the cellular and pathway level. We used primary cell cultures, a new cell line and sections directly derived from freshly resected samples so as to maximize relevance to the pathology. We provide evidence for the existence of two main non-overlapping populations in these tumours that can be easily distinguished on the basis of differential expression of OLIG1 and SOX9. We used double-stainings to further characterize these cells and found that SOX9^+^ cells are related to the astrocytic lineage while OLIG1^+^ cells are more associated with the oligodendrocyte lineage. The phenotype of these two cell types is summarized on Fig. 6F and differs in several aspects. At the transcriptional level, astro-like cells express high levels of HEY1/2, ID4 and SOX9 while oligo-like cells display strong stainings for ASCL1/MASH1, OLIG1 and SOX4/SOX8. We also identified that oligo-like cells had higher levels of 3 receptors (EGFR, PDGFRA and PTPRZ1) than astro-like cells. Finally, we observed a higher level of APOE, BMP4, KCNN3/SK3 and of CRYAB proteins in astro-like cells whereas the IDH1 enzyme appears to be more expressed in oligo-like cells in oligodendroglioma. SOX9^+^ and OLIG1^+^ cells were observed in 6/6 of the samples we explored suggesting that they are likely to be present in a majority of DLGG with IDH1 R132H mutation. These 2 cell types account for over 50% of tumoural cells but their relative proportion is variable between patients (Fig. 1A and S2A).

Former studies based on electron microscopy [46] and immunohistochemistry of DLGG [47–49] have described the presence of a phenotypic heterogeneity within tumoural cells. In particular, in oligodendrogliomas, subpopulations of gliofibrillary oligodendrocytes have been reported [50–52]. While this work was performed, the cellular composition of IDH1-mutated DLGG was also explored using the recent technological approach based on single cell RNA seq [21,22]. These studies reported the presence of three subpopulations of malignant cells: non-proliferating cells differentiated along the astrocytic and oligodendrocytic lineages and proliferative undifferentiated cells that resemble neural stem/progenitor cells. Bioinformatics analysis further established the formation of a tumoural lineage in DLGG and enrichment for SOX4 expression in the proliferative undifferentiated subpopulation. Here we found a widespread expression of SOX4 in the tumor, in particular in OLIG1^+^ cells (Fig. 2C, D and S4C, D) and so far we could not clearly assign this marker to a specific subpopulation acting as a cancer stem cells. Collectively, the present immunofluorescence analysis, former electron microscopic/IHC studies and single cell RNA sequencing approaches all converge to identify the presence of astro-like and oligo-like cells in DLGG. In the normal brain, oligodendrocytes and astrocytes have very distinct properties. For instance, they differ at the metabolic, angiogenic and migration levels. It remains to be established whether oligo-like and astro-like cells found in DLGG show similar dichotomic properties.

One unexpected finding is the highest expression of IDH1 enzyme in oligo-like cells than in astro-like cells in oligodendroglioma. This was observed using an antibody for the mutated form of IDH1 (R132H) but also confirmed with an antibody against wild type IDH1. In fact, in accordance with these findings, single cell RNA databases for human and mouse adult brain cells revealed that IDH1 is more expressed in the oligodendrocytic lineage, especially in immature oligodendrocytes, than in the astrocytic lineage (Fig. S9). We also found that IDH1 expression is also negatively regulated by activation of Notch1 signalling (Fig. 5A, B) which repress several oligodendrocytic genes in DLGG cells. What could be the function of IDH1 in OLIG1^+^ tumoural cells? Compared to IDH2 that is located in the mitochondria to generate alpha-ketoglutarate and NADH, H^+^ in Krebs cycle, IDH1 enzyme is located in the cytoplasm where it generates alpha-ketoglutarate and NADPH, H^+^. NADPH, H^+^ is a key metabolite for lipid and myelin synthesis [53] and thus the high expression of IDH1 observed in the oligodendrocyte lineage may serve the requirement for a high lipid metabolism in these cells.

Another unexpected finding is the specific expression of CRYAB in astro-like tumoural cells. This crystallin protein is typically expressed in inflammatory contexts such as multiple sclerosis [54]. This small heat-shock protein works as an assembly chaperon and have anti-apoptotic, neuroprotective and anti-inflammatory functions [55]. It can accumulate in large amounts in astrocytes in pathological conditions [56]. In cancer, CRYAB can act as an oncoprotein or a tumor suppressor [55]. CRYAB is a transcriptional target of BMP signalling in endothelial cells [57] and accordingly, we found it to be strongly upregulated by BMP signalling (Fig. 6 C-E). CRYAB is also upregulated by NICD overexpression in LGG275 cells (Fig. 5B) suggesting a potential double regulation by Notch1 and BMP signallings. A potential link between CRYAB, Notch1 and BMP pathways is also supported by DLGG RNA profiles in TCGA database showing a significant correlation between CRYAB, BMP2 and HEY2 (Fig S14A). The role of CRYAB in DLGG cells is not known and remains to be elucidated.

The discovery of astro- and oligo-like tumoural cells prompted us to analyze the expression of the G protein-coupled receptor GPR17. In the normal human and mouse brain, GPR17 is almost exclusively expressed by oligodendrocyte progenitors and committed oligodendrocytic cells [58]. It participates in the differentiation and maturation process of oligodendrocytes by binding to several ligands such as leukotrienes and uracil nucleotides. Considering its expression profile, we expected GPR17 to be only expressed in OLIG1^+^ cells however we found it to be also present in a substantial number of SOX9^+^ astrocytic cells. In an inflammatory context, GPR17 can be expressed in non-oligodendrocytic cells [58] so a similar phenomenon could occur in DLGG. Alternatively, astro-like tumoural cells could be derived from the differentiation of GPR17^+^ oligo-like tumoural cells and maintain this marker expression. Whatever its origin and considering that GPR17 activation can impede progenitor differentiation [58], its expression in astro- and oligo-like cells may contribute to the differentiation blockage observed in DLGG cells [8].

Using DLGG primary cultures and a new IDH1/ATRX mutated cell line, we found that the Notch1 pathway deeply influences DLGG cell phenotype. Indeed, by manipulating this pathway through different approaches (constitutively-activated form, pharmacology and DLL4 ligand), we found that Notch1 activation globally reduced the expression of oligodendrocytic genes, while concomitantly increasing KCNN3 and APOE, which are typically associated with the astrocytic lineage. This was accompanied by reduction of proliferation in LGG275 cells as previously seen in high grade gliomas [27,59]. Using non-tumoural human O4-purified cells, we also found that activation of Notch1 reduced oligodendrocytic gene expression while upregulating CRYAB, KCNN3 and SOX9. These results are consistent with the literature showing that Notch1 activation in neural precursor cells promotes astroglial lineage entry [60] and block oligodendrocyte differentiation [61]. Considering the high expression of SOX9 in DLGG astro-like cells (Fig. 1A and S2A) and that SOX9 is a target of Notch1 during nervous system development [44,62] and in other non-neural tissues [63,64], we also expected SOX9 expression to be upregulated by Notch1 activation in DLGG primary cultures and in the LGG275 cell line. This was however not observed in these cells which in fact already expressed a high and may be saturated level of SOX9 (Fig. S10C, S12B). This may result from EGF receptor activation by EGF in the media that can upregulate SOX9 as seen in other cell types [65]. In contrast, in non-tumoural oligodendrocytic O4^+^ cells, which do not express SOX9 (Fig. S11C), both SOX9 gene and protein were strongly induced by Notch1 activation. To our knowledge, this is the first demonstration of a Notch1-induced SOX9 expression in human oligodendrocytic cells. Besides modifying their phenotype, we also found that Notch1 activation modifies the electrophysiological properties of DLGG cells as NICD overexpression induced the expression of electrophysiologically active KCNN3/SK3 channels. Considering the high fold change (between 20 to 100) in KCNN3 expression induced by Notch1 activation in the 3 explored cell types, it is possible that this channel is a direct target of this pathway. Ca2^+^-activated K^+^ channels are involved in membrane hyperpolarization but also regulate cellular shape, migration and proliferation [66]. Likewise, they may also regulate these properties in DLGG cells. Last, we found that Notch1 activation reduces cell proliferation in the cell line we isolated. This result is consistent with the work by Giachino et al [67] showing that in a PDGF-driven mouse glioma model, genetic activation of the Notch1 pathway reduces glioma growth and increases survival. Additionally, in oligodendrogliomas, HEY1 and HEY2 showed reduced expression with advanced disease. Tumours with low HEY2 expression had greater cell density and were more proliferative [68]. Finally, in the TCGA database, expression of HEY1 and HEY2 are reduced in grade III and IV gliomas compared to grade II tumours (Fig. S14B). Collectively, these data fuel the emerging notion that Notch1 pathway acts as a tumor suppressor in DLGG [59].

Despite the presence of a high expression of BMP proteins in DLGG (Fig. S7C), especially in astro-like cells for BMP4 (Fig. 3B and S5B), the effect of BMP2/4 treatment on the genes we explored was less prominent than for Notch1 activation. Only induction of CRYAB was strongly upregulated. We also found a strong induction of NOGGIN by BMP proteins probably reflecting the presence of a negative retro control for this pathway in DLGG cells. With regards to CRYAB, BMPs protect endothelial cells from apoptosis, in part, by upregulating CRYAB that acts as an anti-apoptotic factor [57] and a similar situation could occur in DLGG cells.

## 5. Conclusions

In conclusion, the identification of astro-like SOX9^+^ and oligo-like OLIG1^+^ tumoural cells in DLGG raise several new questions. Do these cells show the same sensitivity to treatment? Do they have the same ability to invade the brain, a major obstacle to treat these tumours? Can they interconvert in the tumor? These issues merit further investigation to derive new therapeutic strategies against these tumours.

## Supporting information

Supplementary Informations

## Supplementary Materials

Figure S1: Preferential astrocytic and oligodendrocytic expression of markers selected in the study; Figure S2: Two non-overlapping cell subpopulations detected in DLGG; Figure S3: SOX9^+^ cells show specific protein expression; Figure S4: OLIG1^+^ cells express proteins associated to the oligodendrocytic lineage and neural precursor markers; Figure S5: SOX9^+^ cells specifically express various signaling molecules and receptors; Figure S6: Expression profiles for BMP2/4 and NOGGIN mRNA in DLGG databases; Figure S7: Expression of BMP2/4 and NOGGIN in DLGG; Figure S8: OLIG1^+^ cells specifically express/activate various receptors and IDH1; Figure S9: Cell type specific expression of IDH1 in adult mouse and human brain; Figure S10: Phenotypic characterization of tumoral O4^+^ cells isolated from DLGG resections; Figure S11: Phenotypic characterization of non-tumoral O4^+^ cells isolated from DLGG samples; Figure S12: Phenotypic characterization of the LGG275 cell line. Figure S13: DLL4 effect on cell phenotype and regulation of SOX9 by Notch1 signaling; Figure S14: Database mining for HEY1/2 expression and correlation in different glioma grades. Table S1: Detailed information of the patients used in the article; Table S2: List of cellular markers and supporting references for their specific expression; Table S3: PCR primer pairs used for quantitative RT-PCR; Table S4: Antibodies used for immunostaining.

## Author Contributions

Conceptualization, Meera Augustus and Jean-philippe Hugnot; Data curation, Meera Augustus and Jean-philippe Hugnot; Formal analysis, Meera Augustus, Donovan Pineau, Franck Aimond, Frederique Scamps, William Ritchie, Hugues Duffau and Jean-philippe Hugnot; Funding acquisition, Meera Augustus and Jean-philippe Hugnot; Investigation, Meera Augustus, Donovan Pineau, Franck Aimond, Safa Azar, Nicolas Leventoux, Frederique Scamps, Sophie Muxel, Amelie Darlix, Catherine Goze, Valerie Rigau, Hugues Duffau and Jean-philippe Hugnot; Methodology, Meera Augustus, Franck Aimond, Frederique Scamps and Jean-philippe Hugnot; Project administration, Jean-philippe Hugnot; Resources, David Lecca and Hugues Duffau; Software, William Ritchie; Supervision, Jean-philippe Hugnot; Validation, Meera Augustus, Donovan Pineau, Franck Aimond, Safa Azar, Nicolas Leventoux, Frederique Scamps, Amelie Darlix, William Ritchie, Catherine Goze, Valerie Rigau, Hugues Duffau and Jean-philippe Hugnot; Visualization, Meera Augustus, Sophie Muxel and Jean-philippe Hugnot; Writing – original draft, Meera Augustus and Jean-philippe Hugnot; Writing – review & editing, Meera Augustus, Donovan Pineau and Jean-philippe Hugnot.

## Institutional Review Board Statement

Tumour samples were obtained from patients with written consents from the “Centre de Ressources Biologiques” located at the Montpellier hospital (information for patients are listed in Table S1) with agreements n° IRB-MTP_2020_09_202000583 and 2019-IRB-MTP-10-15 obtained from the Montpellier hospital Institutional Review Board (IRB ID: 202000583, 198711). The study was conducted according to the guidelines of the Declaration of Helsinki, and approved by the Institutional Review Board of CHU Hospital of Montpellier (202000583, n° 198711) (protocol codes: RECH/P487 (17/10/2019) and 2020000583 (17/9/2020)).

## Informed Consent Statement

Informed consent was obtained from all subjects involved in the study.

## Acknowledgments

We thank Amélie Sarrazin and Christophe Duperray from MRI RIO biocampus facilities for their technical assistance as well as RHEM facility and CHU of Montpellier, Centre de Ressources Biologiques du CHU de Montpellier (CRB), Collection Neurologie, F-34285 Montpellier, France. We thank A. Monteil and C. Lemmers from the Vectorology facility, PVM, Biocampus Montpellier, CNRS UMS3426 for producing lentiviruses used in the study.

## References

1. Ostrom, Q.T.; Gittleman, H.; Truitt, G.; Boscia, A.; Kruchko, C.; Barnholtz-Sloan, J.S. CBTRUS Statistical Report: Primary Brain and Other Central Nervous System Tumors Diagnosed in the United States in 2011-2015. Neuro Oncol 2018, 20, iv1–iv86, doi:10.1093/neuonc/noy131.

2. Louis, D.N.; Ohgaki, H.; Wiestler, O.D.; Cavenee, W.K.; Burger, P.C.; Jouvet, A.; Scheithauer, B.W.; Kleihues, P. The 2007 WHO classification of tumours of the central nervous system. Acta Neuropathol 2007, 114, 97–109, doi:10.1007/s00401-007-0243-4.

3. Pekmezci, M.; Rice, T.; Molinaro, A.M.; Walsh, K.M.; Decker, P.A.; Hansen, H.; Sicotte, H.; Kollmeyer, T.M.; McCoy, L.S.; Sarkar, G., et al. Adult infiltrating gliomas with WHO 2016 integrated diagnosis: additional prognostic roles of ATRX and TERT. Acta Neuropathol 2017, 133, 1001–1016, doi:10.1007/s00401-017-1690-1.

4. Mandonnet, E.; Delattre, J.Y.; Tanguy, M.L.; Swanson, K.R.; Carpentier, A.F.; Duffau, H.; Cornu, P.; Van Effenterre, R.; Alvord, E.C., Jr.; Capelle, L. Continuous growth of mean tumor diameter in a subset of grade II gliomas. Ann Neurol 2003, 53, 524–528, doi:10.1002/ana.10528.

5. Jaeckle, K.A.; Decker, P.A.; Ballman, K.V.; Flynn, P.J.; Giannini, C.; Scheithauer, B.W.; Jenkins, R.B.; Buckner, J.C. Transformation of low grade glioma and correlation with outcome: an NCCTG database analysis. J Neurooncol 2011, 104, 253–259, doi:10.1007/s11060-010-0476-2.

6. Oberheim Bush, N.A.; Chang, S. Treatment Strategies for Low-Grade Glioma in Adults. J Oncol Pract 2016, 12, 1235–1241, doi:10.1200/JOP.2016.018622.

7. Parsons, D.W.; Jones, S.; Zhang, X.; Lin, J.C.; Leary, R.J.; Angenendt, P.; Mankoo, P.; Carter, H.; Siu, I.M.; Gallia, G.L., et al. An integrated genomic analysis of human glioblastoma multiforme. Science 2008, 321, 1807–1812, doi:10.1126/science.1164382.

8. Lu, C.; Ward, P.S.; Kapoor, G.S.; Rohle, D.; Turcan, S.; Abdel-Wahab, O.; Edwards, C.R.; Khanin, R.; Figueroa, M.E.; Melnick, A., et al. IDH mutation impairs histone demethylation and results in a block to cell differentiation. Nature 2012, 483, 474–478, doi:10.1038/nature10860.

9. Turcan, S.; Rohle, D.; Goenka, A.; Walsh, L.A.; Fang, F.; Yilmaz, E.; Campos, C.; Fabius, A.W.; Lu, C.; Ward, P.S., et al. IDH1 mutation is sufficient to establish the glioma hypermethylator phenotype. Nature 2012, 483, 479–483, doi:10.1038/nature10866.

10. Louis, D.N.; Perry, A.; Reifenberger, G.; von Deimling, A.; Figarella-Branger, D.; Cavenee, W.K.; Ohgaki, H.; Wiestler, O.D.; Kleihues, P.; Ellison, D.W. The 2016 World Health Organization Classification of Tumors of the Central Nervous System: a summary. Acta Neuropathol 2016, 131, 803–820, doi:10.1007/s00401-016-1545-1.

11. Cahill, D.P.; Louis, D.N.; Cairncross, J.G. Molecular background of oligodendroglioma: 1p/19q, IDH, TERT, CIC and FUBP1. CNS Oncol 2015, 4, 287–294, doi:10.2217/cns.15.32.

12. Cancer Genome Atlas Research, N.; Brat, D.J.; Verhaak, R.G.; Aldape, K.D.; Yung, W.K.; Salama, S.R.; Cooper, L.A.; Rheinbay, E.; Miller, C.R.; Vitucci, M., et al. Comprehensive, Integrative Genomic Analysis of Diffuse Lower-Grade Gliomas. N Engl J Med 2015, 372, 2481–2498, doi:10.1056/NEJMoa1402121.

13. Jiao, Y.; Killela, P.J.; Reitman, Z.J.; Rasheed, A.B.; Heaphy, C.M.; de Wilde, R.F.; Rodriguez, F.J.; Rosemberg, S.; Oba-Shinjo, S.M.; Nagahashi Marie, S.K., et al. Frequent ATRX, CIC, FUBP1 and IDH1 mutations refine the classification of malignant gliomas. Oncotarget 2012, 3, 709–722, doi:10.18632/oncotarget.588.

14. Furnari, F.B.; Cloughesy, T.F.; Cavenee, W.K.; Mischel, P.S. Heterogeneity of epidermal growth factor receptor signalling networks in glioblastoma. Nat Rev Cancer 2015, 15, 302–310, doi:10.1038/nrc3918.

15. Szerlip, N.J.; Pedraza, A.; Chakravarty, D.; Azim, M.; McGuire, J.; Fang, Y.; Ozawa, T.; Holland, E.C.; Huse, J.T.; Jhanwar, S., et al. Intratumoral heterogeneity of receptor tyrosine kinases EGFR and PDGFRA amplification in glioblastoma defines subpopulations with distinct growth factor response. Proc Natl Acad Sci U S A 2012, 109, 3041–3046, doi:10.1073/pnas.1114033109.

16. Lino, M.M.; Merlo, A. PI3Kinase signaling in glioblastoma. J Neurooncol 2011, 103, 417–427, doi:10.1007/s11060-010-0442-z.

17. Pierscianek, D.; Kim, Y.H.; Motomura, K.; Mittelbronn, M.; Paulus, W.; Brokinkel, B.; Keyvani, K.; Wrede, K.; Nakazato, Y.; Tanaka, Y., et al. MET gain in diffuse astrocytomas is associated with poorer outcome. Brain Pathol 2013, 23, 13–18, doi:10.1111/j.1750-3639.2012.00609.x.

18. Motomura, K.; Mittelbronn, M.; Paulus, W.; Brokinkel, B.; Keyvani, K.; Sure, U.; Wrede, K.; Nakazato, Y.; Tanaka, Y.; Nonoguchi, N., et al. PDGFRA gain in low-grade diffuse gliomas. J Neuropathol Exp Neurol 2013, 72, 61–66, doi:10.1097/NEN.0b013e31827c4b5b.

19. Bao, Z.S.; Chen, H.M.; Yang, M.Y.; Zhang, C.B.; Yu, K.; Ye, W.L.; Hu, B.Q.; Yan, W.; Zhang, W.; Akers, J., et al. RNA-seq of 272 gliomas revealed a novel, recurrent PTPRZ1-MET fusion transcript in secondary glioblastomas. Genome Res 2014, 24, 1765–1773, doi:10.1101/gr.165126.113.

20. Azar, S.; Leventoux, N.; Ripoll, C.; Rigau, V.; Goze, C.; Lorcy, F.; Bauchet, L.; Duffau, H.; Guichet, P.O.; Rothhut, B., et al. Cellular and molecular characterization of IDH1-mutated diffuse low grade gliomas reveals tumor heterogeneity and absence of EGFR/PDGFRalpha activation. Glia 2018, 66, 239–255, doi:10.1002/glia.23240.

21. Tirosh, I.; Venteicher, A.S.; Hebert, C.; Escalante, L.E.; Patel, A.P.; Yizhak, K.; Fisher, J.M.; Rodman, C.; Mount, C.; Filbin, M.G., et al. Single-cell RNA-seq supports a developmental hierarchy in human oligodendroglioma. Nature 2016, 539, 309–313, doi:10.1038/nature20123.

22. Venteicher, A.S.; Tirosh, I.; Hebert, C.; Yizhak, K.; Neftel, C.; Filbin, M.G.; Hovestadt, V.; Escalante, L.E.; Shaw, M.L.; Rodman, C., et al. Decoupling genetics, lineages, and microenvironment in IDH-mutant gliomas by single-cell RNA-seq. Science 2017, 355, doi:10.1126/science.aai8478.

23. Leventoux, N.; Augustus, M.; Azar, S.; Riquier, S.; Villemin, J.P.; Guelfi, S.; Falha, L.; Bauchet, L.; Goze, C.; Ritchie, W., et al. Transformation Foci in IDH1-mutated Gliomas Show STAT3 Phosphorylation and Downregulate the Metabolic Enzyme ETNPPL, a Negative Regulator of Glioma Growth. Sci Rep 2020, 10, 5504, doi:10.1038/s41598-020-62145-1.

24. Goze, C.; Bezzina, C.; Goze, E.; Rigau, V.; Maudelonde, T.; Bauchet, L.; Duffau, H. 1P19Q loss but not IDH1 mutations influences WHO grade II gliomas spontaneous growth. J Neurooncol 2012, 108, 69–75, doi:10.1007/s11060-012-0831-6.

25. Engin, F.; Yao, Z.; Yang, T.; Zhou, G.; Bertin, T.; Jiang, M.M.; Chen, Y.; Wang, L.; Zheng, H.; Sutton, R.E., et al. Dimorphic effects of Notch signaling in bone homeostasis. Nat Med 2008, 14, 299–305, doi:10.1038/nm1712.

26. Avliyakulov, N.K.; Rajavel, K.S.; Le, K.M.; Guo, L.; Mirsadraei, L.; Yong, W.H.; Liau, L.M.; Li, S.; Lai, A.; Nghiemphu, P.L., et al. C-terminally truncated form of alphaB-crystallin is associated with IDH1 R132H mutation in anaplastic astrocytoma. J Neurooncol 2014, 117, 53–65, doi:10.1007/s11060-014-1371-z.

27. Guichet, P.O.; Guelfi, S.; Teigell, M.; Hoppe, L.; Bakalara, N.; Bauchet, L.; Duffau, H.; Lamszus, K.; Rothhut, B.; Hugnot, J.P. Notch1 stimulation induces a vascularization switch with pericyte-like cell differentiation of glioblastoma stem cells. Stem Cells 2015, 33, 21–34, doi:10.1002/stem.1767.

28. Gross, R.E.; Mehler, M.F.; Mabie, P.C.; Zang, Z.; Santschi, L.; Kessler, J.A. Bone morphogenetic proteins promote astroglial lineage commitment by mammalian subventricular zone progenitor cells. Neuron 1996, 17, 595–606, doi:10.1016/s0896-6273(00)80193-2.

29. Srikanth, M.; Kim, J.; Das, S.; Kessler, J.A. BMP signaling induces astrocytic differentiation of clinically derived oligodendroglioma propagating cells. Mol Cancer Res 2014, 12, 283–294, doi:10.1158/1541-7786.MCR-13-0349.

30. Ekstrand, A.J.; Sugawa, N.; James, C.D.; Collins, V.P. Amplified and rearranged epidermal growth factor receptor genes in human glioblastomas reveal deletions of sequences encoding portions of the N- and/or C-terminal tails. Proc Natl Acad Sci U S A 1992, 89, 4309–4313, doi:10.1073/pnas.89.10.4309.

31. Huang, P.H.; Xu, A.M.; White, F.M. Oncogenic EGFR signaling networks in glioma. Sci Signal 2009, 2, re6, doi:10.1126/scisignal.287re6.

32. Muller, S.; Kunkel, P.; Lamszus, K.; Ulbricht, U.; Lorente, G.A.; Nelson, A.M.; von Schack, D.; Chin, D.J.; Lohr, S.C.; Westphal, M., et al. A role for receptor tyrosine phosphatase zeta in glioma cell migration. Oncogene 2003, 22, 6661–6668, doi:10.1038/sj.onc.1206763.

33. Ohgaki, H.; Dessen, P.; Jourde, B.; Horstmann, S.; Nishikawa, T.; Di Patre, P.L.; Burkhard, C.; Schuler, D.; Probst-Hensch, N.M.; Maiorka, P.C., et al. Genetic pathways to glioblastoma: a population-based study. Cancer Res 2004, 64, 6892–6899, doi:10.1158/0008-5472.CAN-04-1337.

34. Pedeutour-Braccini, Z.; Burel-Vandenbos, F.; Goze, C.; Roger, C.; Bazin, A.; Costes-Martineau, V.; Duffau, H.; Rigau, V. Microfoci of malignant progression in diffuse low-grade gliomas: towards the creation of an intermediate grade in glioma classification? Virchows Arch 2015, 466, 433–444, doi:10.1007/s00428-014-1712-5.

35. Ulbricht, U.; Brockmann, M.A.; Aigner, A.; Eckerich, C.; Muller, S.; Fillbrandt, R.; Westphal, M.; Lamszus, K. Expression and function of the receptor protein tyrosine phosphatase zeta and its ligand pleiotrophin in human astrocytomas. J Neuropathol Exp Neurol 2003, 62, 1265–1275, doi:10.1093/jnen/62.12.1265.

36. Geisbrecht, B.V.; Gould, S.J. The human PICD gene encodes a cytoplasmic and peroxisomal NADP(+)-dependent isocitrate dehydrogenase. J Biol Chem 1999, 274, 30527–30533, doi:10.1074/jbc.274.43.30527.

37. Trotter, J.; Schachner, M. Cells positive for the O4 surface antigen isolated by cell sorting are able to differentiate into astrocytes or oligodendrocytes. Brain Res Dev Brain Res 1989, 46, 115–122, doi:10.1016/0165-3806(89)90148-x.

38. Yordanova, Y.N.; Duffau, H. Supratotal resection of diffuse gliomas - an overview of its multifaceted implications. Neurochirurgie 2017, 63, 243–249, doi:10.1016/j.neuchi.2016.09.006.

39. Dromard, C.; Bartolami, S.; Deleyrolle, L.; Takebayashi, H.; Ripoll, C.; Simonneau, L.; Prome, S.; Puech, S.; Tran, V.B.; Duperray, C., et al. NG2 and Olig2 expression provides evidence for phenotypic deregulation of cultured central nervous system and peripheral nervous system neural precursor cells. Stem Cells 2007, 25, 340–353, doi:10.1634/stemcells.2005-0556.

40. Ligon, K.L.; Alberta, J.A.; Kho, A.T.; Weiss, J.; Kwaan, M.R.; Nutt, C.L.; Louis, D.N.; Stiles, C.D.; Rowitch, D.H. The oligodendroglial lineage marker OLIG2 is universally expressed in diffuse gliomas. J Neuropathol Exp Neurol 2004, 63, 499–509, doi:10.1093/jnen/63.5.499.

41. Li, J.L.; Sainson, R.C.; Shi, W.; Leek, R.; Harrington, L.S.; Preusser, M.; Biswas, S.; Turley, H.; Heikamp, E.; Hainfellner, J.A., et al. Delta-like 4 Notch ligand regulates tumor angiogenesis, improves tumor vascular function, and promotes tumor growth in vivo. Cancer Res 2007, 67, 11244–11253, doi:10.1158/0008-5472.CAN-07-0969.

42. Mailhos, C.; Modlich, U.; Lewis, J.; Harris, A.; Bicknell, R.; Ish-Horowicz, D. Delta4, an endothelial specific notch ligand expressed at sites of physiological and tumor angiogenesis. Differentiation 2001, 69, 135–144, doi:10.1046/j.1432-0436.2001.690207.x.

43. Adelman, J.P.; Maylie, J.; Sah, P. Small-conductance Ca2+-activated K+ channels: form and function. Annu Rev Physiol 2012, 74, 245–269, doi:10.1146/annurev-physiol-020911-153336.

44. Martini, S.; Bernoth, K.; Main, H.; Ortega, G.D.; Lendahl, U.; Just, U.; Schwanbeck, R. A critical role for Sox9 in notch-induced astrogliogenesis and stem cell maintenance. Stem Cells 2013, 31, 741–751, doi:10.1002/stem.1320.

45. Kluppel, M.; Wrana, J.L. Turning it up a Notch: cross-talk between TGF beta and Notch signaling. Bioessays 2005, 27, 115–118, doi:10.1002/bies.20187.

46. Baloyannis, S. The fine structure of the isomorphic oligodendroglioma. Anticancer Res 1981, 1, 243–248.

47. Liang, Y.; Bollen, A.W.; Nicholas, M.K.; Gupta, N. Id4 and FABP7 are preferentially expressed in cells with astrocytic features in oligodendrogliomas and oligoastrocytomas. BMC Clin Pathol 2005, 5, 6, doi:10.1186/1472-6890-5-6.

48. Rousseau, A.; Nutt, C.L.; Betensky, R.A.; Iafrate, A.J.; Han, M.; Ligon, K.L.; Rowitch, D.H.; Louis, D.N. Expression of oligodendroglial and astrocytic lineage markers in diffuse gliomas: use of YKL-40, ApoE, ASCL1, and NKX2-2. J Neuropathol Exp Neurol 2006, 65, 1149–1156, doi:10.1097/01.jnen.0000248543.90304.2b.

49. de la Monte, S.M. Uniform lineage of oligodendrogliomas. Am J Pathol 1989, 135, 529–540.

50. Herpers, M.J.; Budka, H. Glial fibrillary acidic protein (GFAP) in oligodendroglial tumors: gliofibrillary oligodendroglioma and transitional oligoastrocytoma as subtypes of oligodendroglioma. Acta Neuropathol 1984, 64, 265–272, doi:10.1007/BF00690392.

51. Kros, J.M.; Schouten, W.C.; Janssen, P.J.; van der Kwast, T.H. Proliferation of gemistocytic cells and glial fibrillary acidic protein (GFAP)-positive oligodendroglial cells in gliomas: a MIB-1/GFAP double labeling study. Acta Neuropathol 1996, 91, 99–103, doi:10.1007/s004010050398.

52. Matyja, E.; Taraszewska, A.; Zabek, M. Phenotypic characteristics of GFAP-immunopositive oligodendroglial tumours Part I: immunohistochemical study. Folia Neuropathol 2001, 39, 19–26.

53. Bourre, J.M. Developmental Synthesis of Myelin Lipids: Origin of Fatty Acids—Specific Role of Nutrition. In Developmental Neurobiohgy, New York, 1989; Vol. 12, pp. 111–154.

54. Ousman, S.S.; Tomooka, B.H.; van Noort, J.M.; Wawrousek, E.F.; O’Connor, K.C.; Hafler, D.A.; Sobel, R.A.; Robinson, W.H.; Steinman, L. Protective and therapeutic role for alphaB-crystallin in autoimmune demyelination. Nature 2007, 448, 474–479, doi:10.1038/nature05935.

55. Zhang, J.; Liu, J.; Wu, J.; Li, W.; Chen, Z.; Yang, L. Progression of the role of CRYAB in signaling pathways and cancers. Onco Targets Ther 2019, 12, 4129–4139, doi:10.2147/OTT.S201799.

56. Iwaki, T.; Kume-Iwaki, A.; Liem, R.K.; Goldman, J.E. Alpha B-crystallin is expressed in non-lenticular tissues and accumulates in Alexander’s disease brain. Cell 1989, 57, 71–78, doi:10.1016/0092-8674(89)90173-6.

57. Ciumas, M.; Eyries, M.; Poirier, O.; Maugenre, S.; Dierick, F.; Gambaryan, N.; Montagne, K.; Nadaud, S.; Soubrier, F. Bone morphogenetic proteins protect pulmonary microvascular endothelial cells from apoptosis by upregulating alpha-B-crystallin. Arterioscler Thromb Vasc Biol 2013, 33, 2577–2584, doi:10.1161/ATVBAHA.113.301976.

58. Lecca, D.; Raffaele, S.; Abbracchio, M.P.; Fumagalli, M. Regulation and signaling of the GPR17 receptor in oligodendroglial cells. Glia 2020, 68, 1957–1967, doi:10.1002/glia.23807.

59. Parmigiani, E.; Taylor, V.; Giachino, C. Oncogenic and Tumor-Suppressive Functions of NOTCH Signaling in Glioma. Cells 2020, 9, doi:10.3390/cells9102304.

60. Gaiano, N.; Fishell, G. The role of notch in promoting glial and neural stem cell fates. Annu Rev Neurosci 2002, 25, 471–490, doi:10.1146/annurev.neuro.25.030702.130823.

61. Wang, S.; Sdrulla, A.D.; diSibio, G.; Bush, G.; Nofziger, D.; Hicks, C.; Weinmaster, G.; Barres, B.A. Notch receptor activation inhibits oligodendrocyte differentiation. Neuron 1998, 21, 63–75, doi:10.1016/s0896-6273(00)80515-2.

62. Muto, A.; Iida, A.; Satoh, S.; Watanabe, S. The group E Sox genes Sox8 and Sox9 are regulated by Notch signaling and are required for Muller glial cell development in mouse retina. Exp Eye Res 2009, 89, 549–558, doi:10.1016/j.exer.2009.05.006.

63. Capaccione, K.M.; Hong, X.; Morgan, K.M.; Liu, W.; Bishop, J.M.; Liu, L.; Markert, E.; Deen, M.; Minerowicz, C.; Bertino, J.R., et al. Sox9 mediates Notch1-induced mesenchymal features in lung adenocarcinoma. Oncotarget 2014, 5, 3636–3650, doi:10.18632/oncotarget.1970.

64. Shih, H.P.; Kopp, J.L.; Sandhu, M.; Dubois, C.L.; Seymour, P.A.; Grapin-Botton, A.; Sander, M. A Notch-dependent molecular circuitry initiates pancreatic endocrine and ductal cell differentiation. Development 2012, 139, 2488–2499, doi:10.1242/dev.078634.

65. Ling, S.; Chang, X.; Schultz, L.; Lee, T.K.; Chaux, A.; Marchionni, L.; Netto, G.J.; Sidransky, D.; Berman, D.M. An EGFR-ERK-SOX9 signaling cascade links urothelial development and regeneration to cancer. Cancer Res 2011, 71, 3812–3821, doi:10.1158/0008-5472.CAN-10-3072.

66. Weaver, A.K.; Bomben, V.C.; Sontheimer, H. Expression and function of calcium-activated potassium channels in human glioma cells. Glia 2006, 54, 223–233, doi:10.1002/glia.20364.

67. Giachino, C.; Boulay, J.L.; Ivanek, R.; Alvarado, A.; Tostado, C.; Lugert, S.; Tchorz, J.; Coban, M.; Mariani, L.; Bettler, B., et al. A Tumor Suppressor Function for Notch Signaling in Forebrain Tumor Subtypes. Cancer Cell 2015, 28, 730–742, doi:10.1016/j.ccell.2015.10.008.

68. Halani, S.H.; Yousefi, S.; Velazquez Vega, J.; Rossi, M.R.; Zhao, Z.; Amrollahi, F.; Holder, C.A.; Baxter-Stoltzfus, A.; Eschbacher, J.; Griffith, B., et al. Multi-faceted computational assessment of risk and progression in oligodendroglioma implicates NOTCH and PI3K pathways. NPJ Precis Oncol 2018, 2, 24, doi:10.1038/s41698-018-0067-9.

