## Supplementary Informations for "Identification of CRYAB^+^ KCNN3^+^ SOX9^+^ astro-like and EGFR^+^ PDGFRA^+^ OLIG1^+^ oligo-like tumoral cells in diffuse low-grade gliomas and implication of Notch1 signalling in their genesis"

A

B

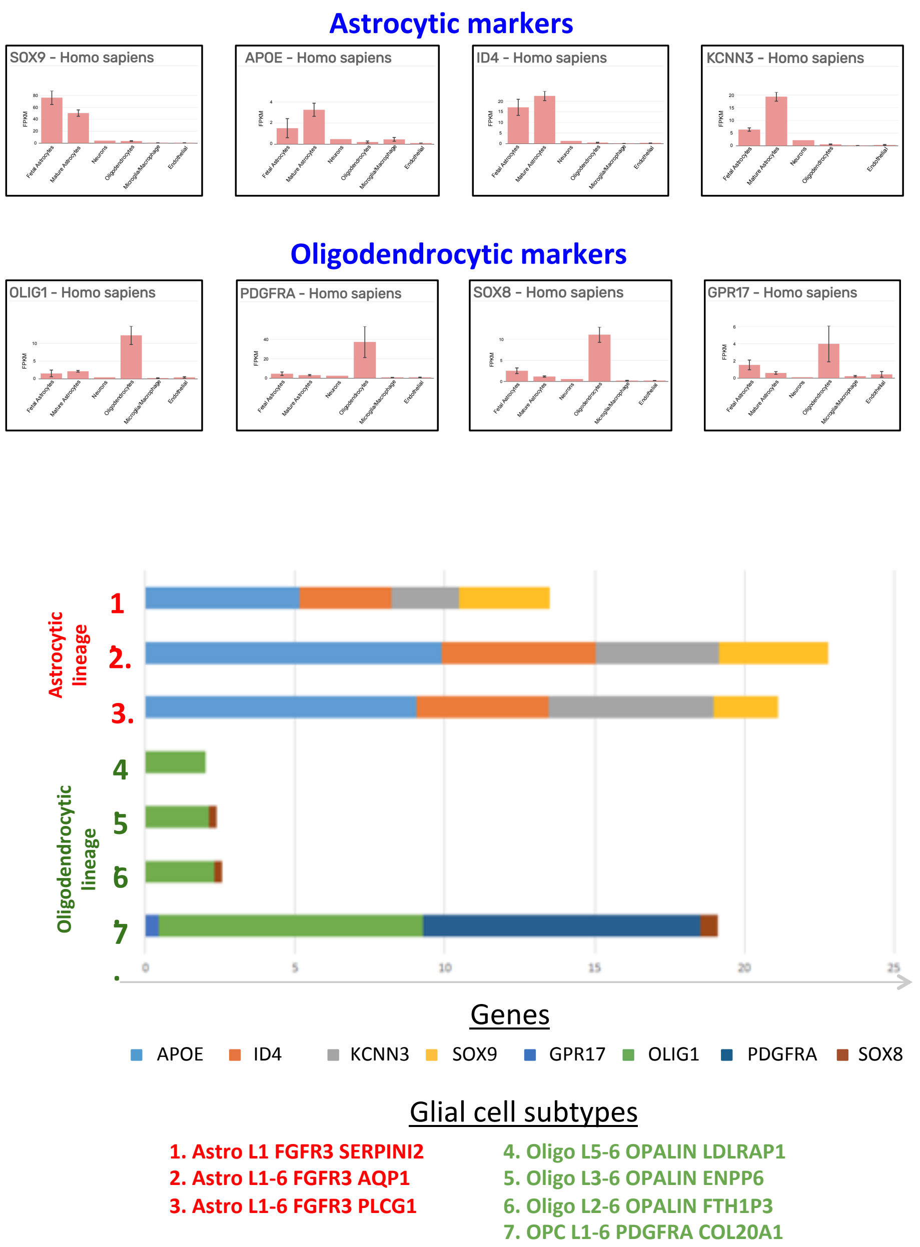


**Figure S1**: Preferential astrocytic and oligodendrocytic expression of markers selected in the study. (A) RNA levels of indicated genes in different human brain cells. Diagrams were retrieved from the brainrnaseq.org database ([1], Dr. Ben Barres's laboratory). (B) RNA expression of indicated genes in different glial human brain cells (M1 cortex) measured by single nuclei RNA seq analysis [2]. This bar chart was drawn using data (expression trimmed mean) extracted from the RNA-Seq Human Cell Types Database 2020 (Allen Institute for Brain Science), available at celltypes.brain-map.org/rnaseq/human_m1_10x. RNA level [2]. Horizontal bars show the level of expression of the 9 indicated genes in the 3 human astrocyte- and 4 oligodendrocyte subtypes defined by the single nuclei RNA seq analysis. APOE, ID4, KCNN3 and SOX9 are preferentially expressed in astrocytic cells while GPR17, OLIG1, PDGFRA and SOX8 are mainly expressed in oligodendrocytic cells.

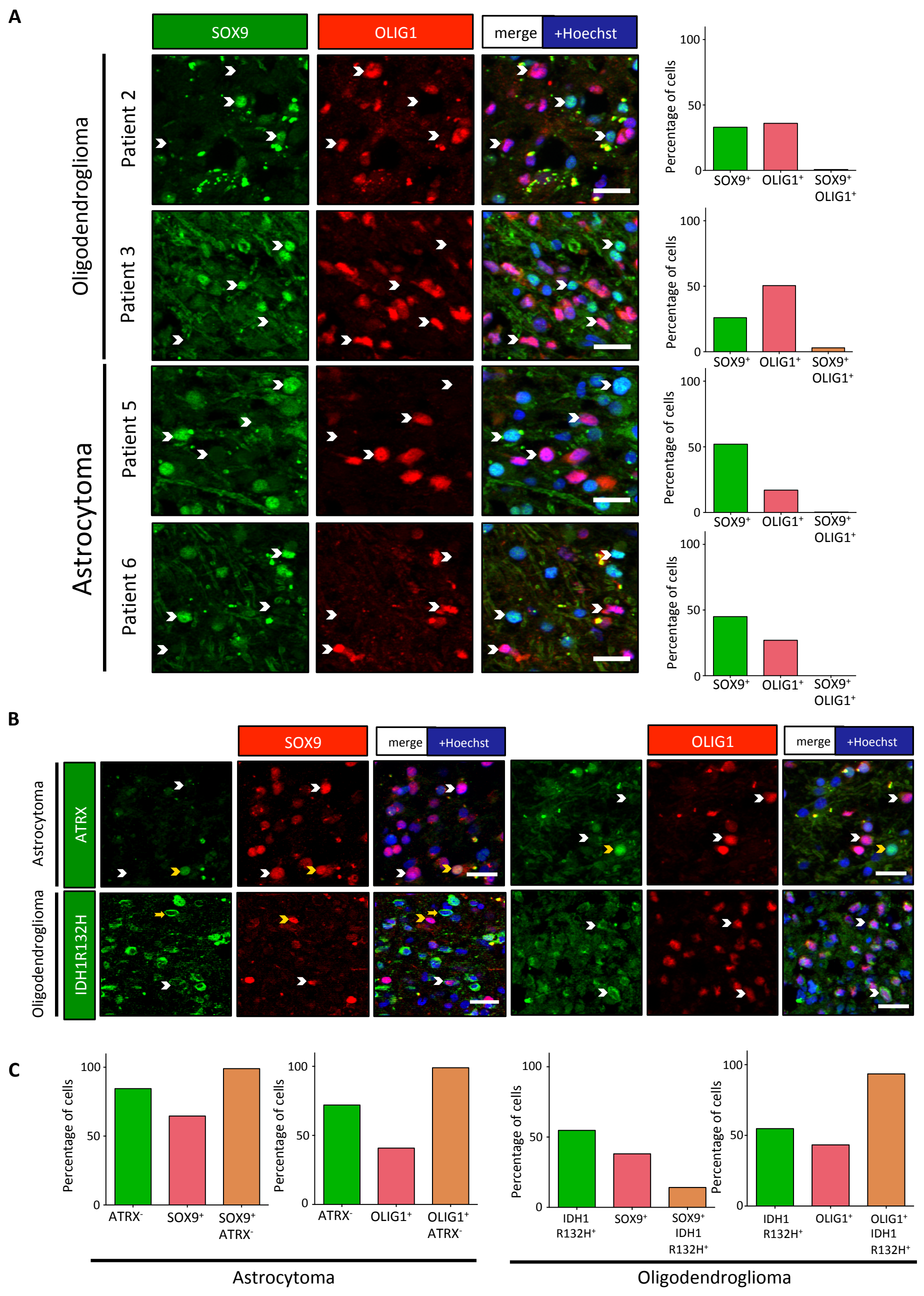


**Figure S2:** Two non-overlapping cell subpopulations detected in DLGG. Double immunofluorescences for indicated markers. (A) Immunofluorescences for SOX9 (green) and OLIG1 (red) on 2 additional oligodendrogliomas and astrocytomas confirmed the presence of two non-overlapping cell subpopulations. White arrowheads mark cells expressing either SOX9 or OLIG1 alone. Scale bars=20 µm. Bar diagrams show the percentage of $SOX9^+$ cells (green), $OLIG1^+$ cells (red) and double positive cells (orange). (B) Tumoral status of $SOX9^+$ and $OLIG1^+$ cells. Upper images: representative images of an astrocytoma section showing the absence of nuclear expression of ATRX in the majority of the $SOX9^+$ and $OLIG1^+$ cells. White arrowheads mark $ATRX^-$ tumoral cells while yellow arrowheads mark non-tumoral cells expressing nuclear ATRX. Lower images: representative images of an oligodendroglioma section double stained for IDH1 R132H protein and SOX9 or OLIG1. White arrowheads mark $OLIG1^+$ and $SOX9^+$ cells expressing IDH1 R132H protein. Yellow arrowheads/arrows show single positive cells. The majority of $OLIG1^+$ cells express IDH1 R132H protein while this mutated protein is only detected in a minor population of $SOX9^+$ cells. Scale bars=20 µm. (C) Quantifications. Left diagrams show the % of $ATRX^-$ cells (green), the % of $SOX9^+$ or $OLIG1^+$ cells (red) and the % of $SOX9^+$ or $OLIG1^+$ cells that are negative for ATRX (orange) in one astrocytoma. Right diagrams show the % of IDH1 $R132H^+$ cells (green), the % of $SOX9^+$ or $OLIG1^+$ cells (red) and the % of $SOX9^+$ or $OLIG1^+$ cells in which the IDH1 R132H mutated protein are detected (orange) in one oligodendroglioma.

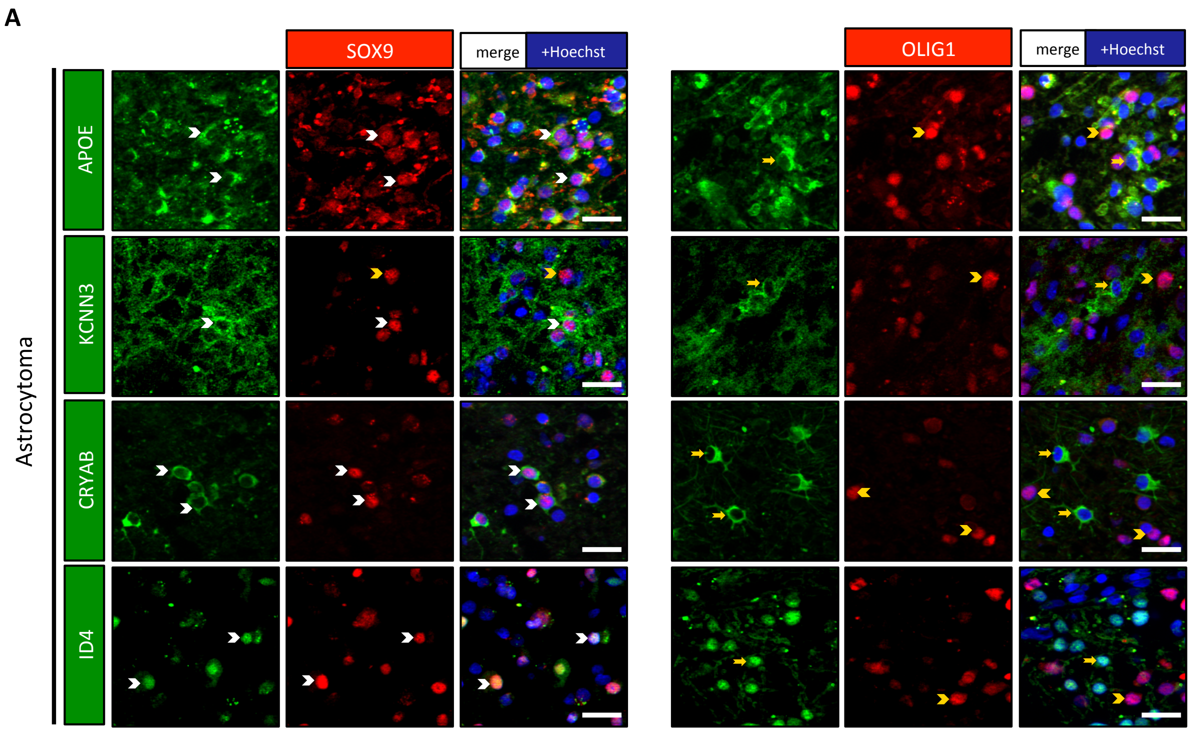


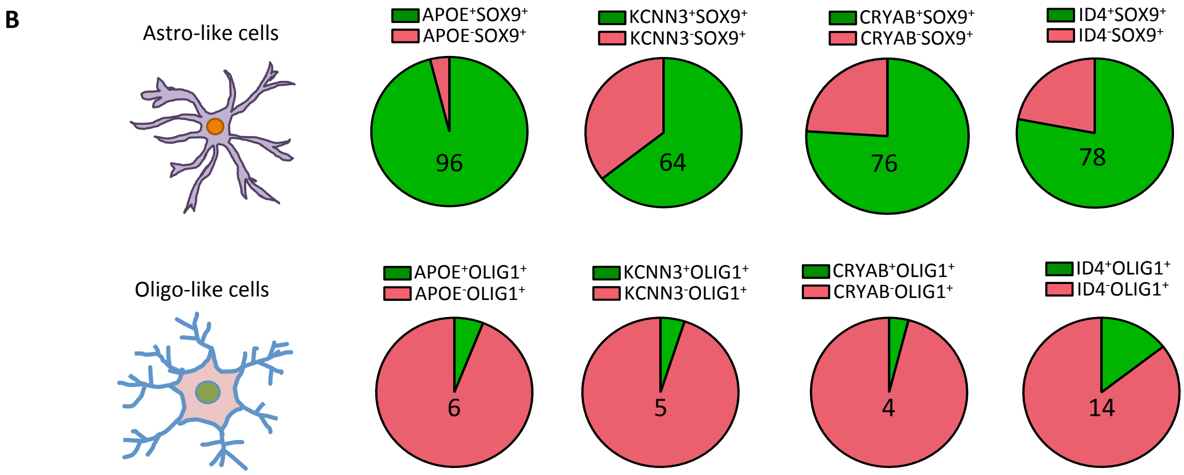


**Figure S3:** $SOX9^{+}$ cells show specific protein expression. Double immunofluorescence for indicated proteins on one astrocytoma. (A) Stainings for SOX9 (left panel) and OLIG1 (right panel) with APOE, CRYAB, KCNN3 and ID4 showed a preferential expression of these proteins in $SOX9^{+}$ cells. White arrowheads show double positive cells while yellow arrowheads/arrows indicate single positive cells. Scale bars=20 µm. (B) Pie diagrams represent the percentage of double positive (green) and single positive (red) cells in astro-like $SOX9^{+}$ (upper lane) and oligo-like $OLIG1^{+}$ (lower lane) cells. Numbers indicate the percentage of double positive cells.

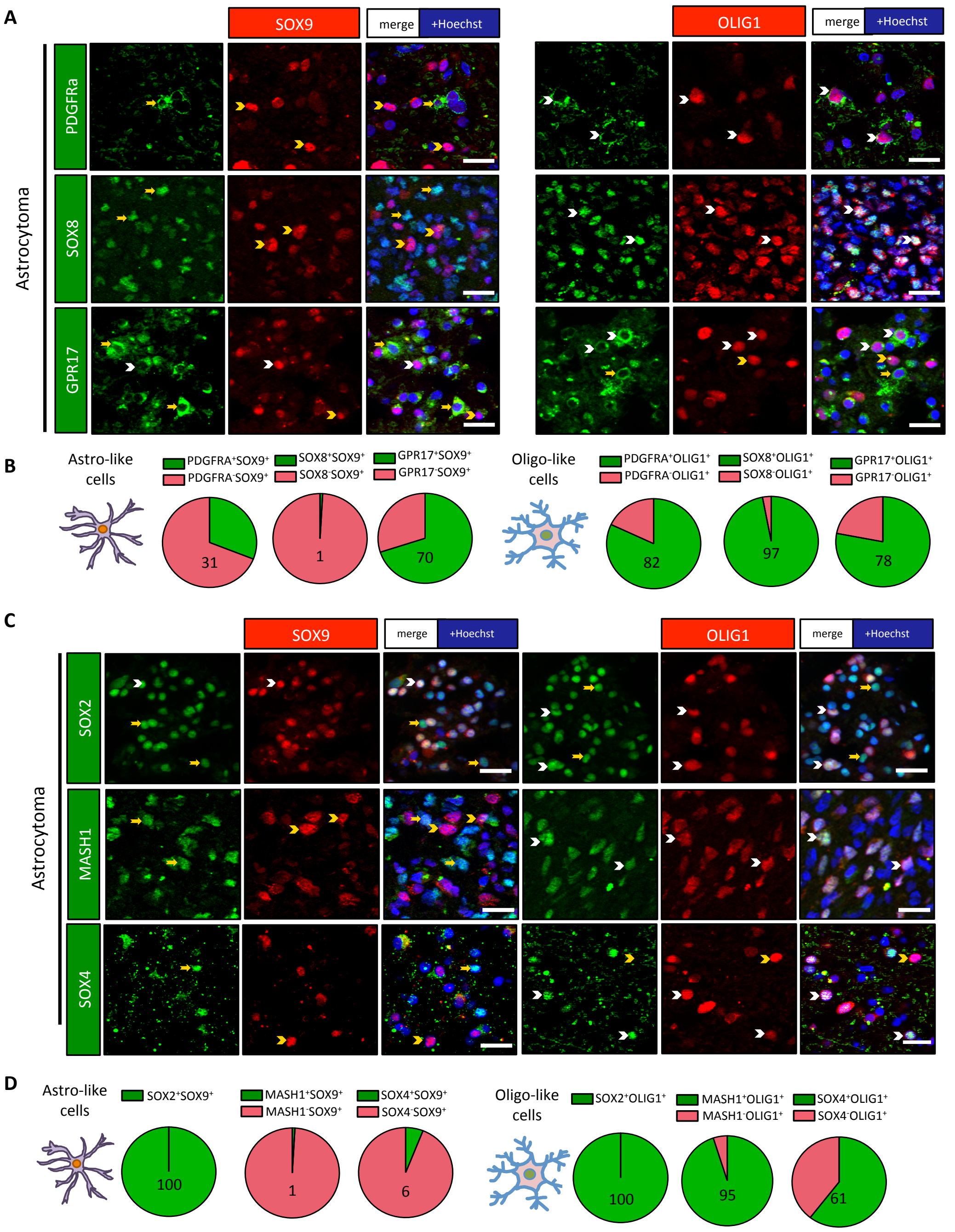


**Figure S4:** OLIG1$^+$ cells express proteins associated to the oligodendrocytic lineage and neural precursor markers. Double immunofluorescences for indicated proteins on one astrocytoma. White arrowheads show double positive cells while yellow arrowheads/arrows show single positive cells. Scale bars=20 µm. (A) Immunofluorescences for PDGFRα and SOX8 revealed their close association with OLIG1$^+$ cells while GPR17 was expressed by both SOX9$^+$ and OLIG1$^+$ cells. (B, D) Pie diagrams represent the percentage of double positive (green) and single positive (red) cells in astro-like SOX9$^+$ (left) and oligo-like OLIG1$^+$ (right) cell populations. Numbers indicate the percentage of double positive cells. (C) Immunofluorescence for MASH1 and SOX4 show their close association with OLIG1$^+$ cells while SOX2 was expressed by both SOX9$^+$ and OLIG1$^+$ cells.

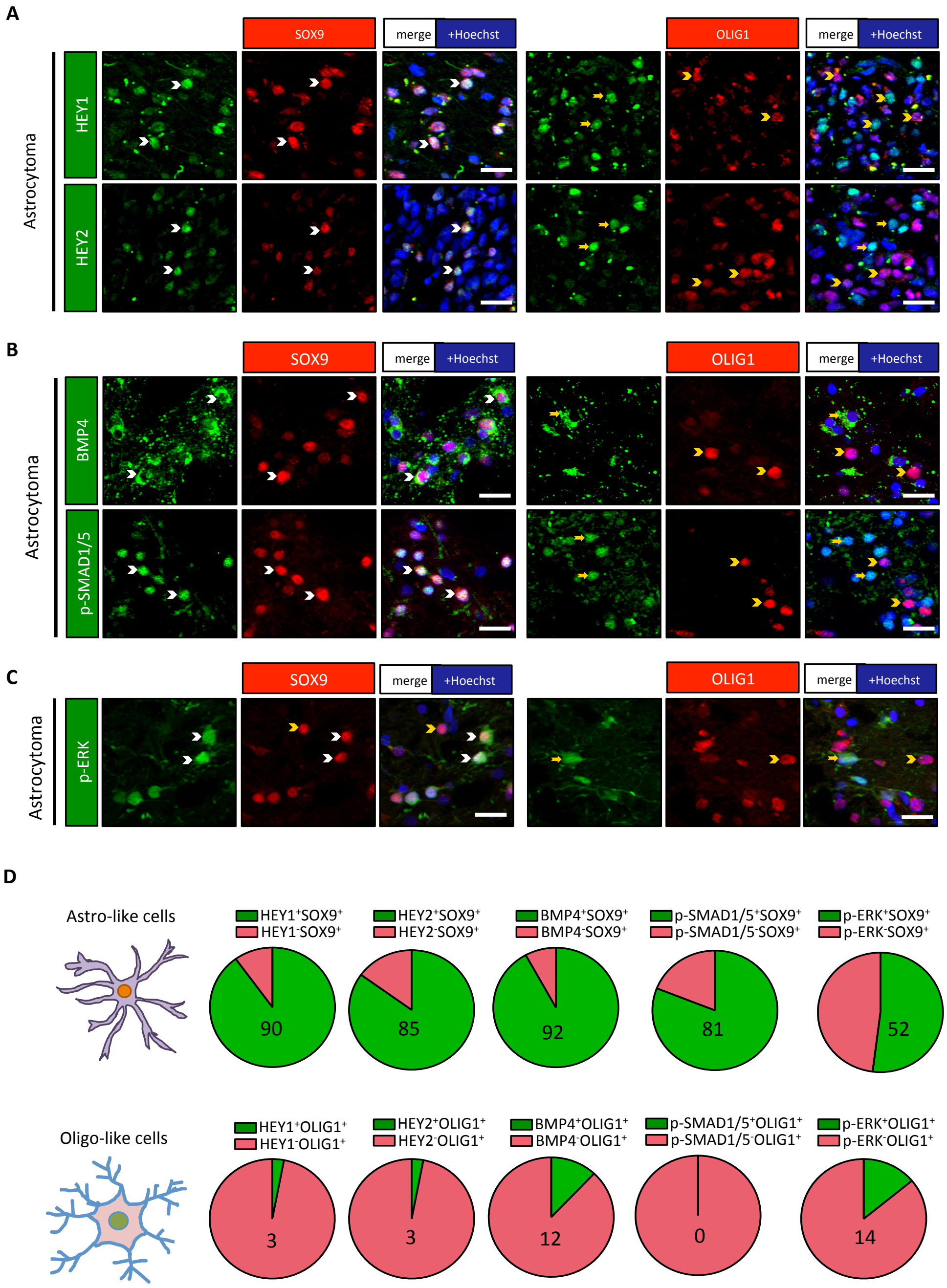


**Figure S5:** SOX9$^+$ cells specifically express various signalling molecules and receptors. Double immunofluorescences for indicated proteins on one astrocytoma. White arrowheads show double positive cells while yellow arrowheads/arrows show single positive cells. Scale bars=20 µm. (A) Stainings for HEY1 or HEY2 with SOX9 and OLIG1 show their specific expression by SOX9$^+$ cells. (B) Stainings for BMP4 and p-SMAD1/5 revealed their specific expression in SOX9$^+$ cells. (C) Stainings for p-ERK show preferential expression in SOX9$^+$ cells. (D) Pie diagrams represent the percentage of double positive (green) and single positive (red) cells in astro-like SOX9$^+$ (upper lane) and oligo-like OLIG1$^+$ (lower lane) cell populations. Numbers indicate the percentage of double positive cells.

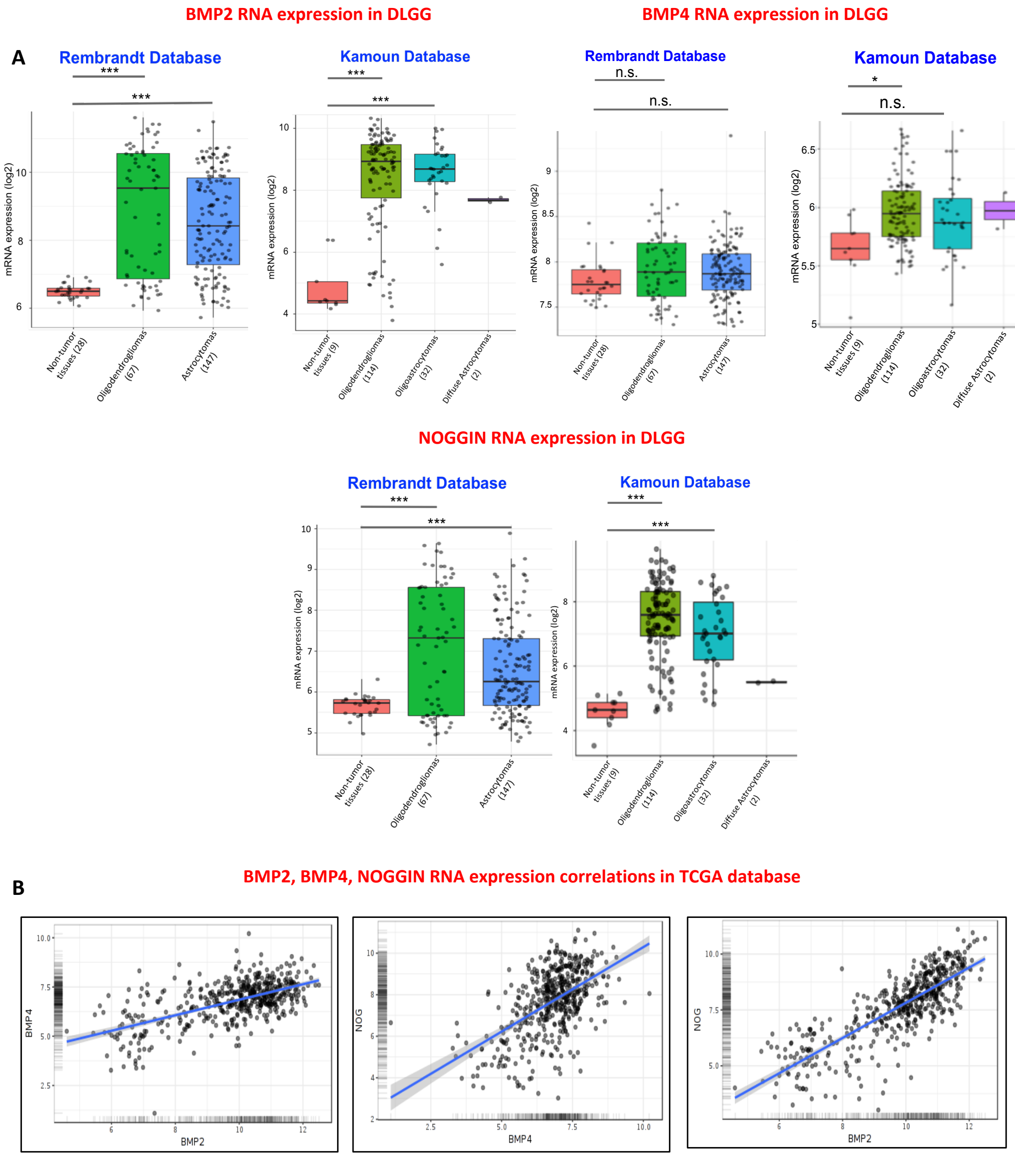


**Correlation matrix (LGG-TCGA database)**
**(Pearson, n=511 cases)**

| | BMP2 | BMP4 | NOGGIN |
|---|---|---|---|
| **BMP2** | | 0.60 | 0.81 |
| **BMP4** | 0.60 | | 0.55 |
| **NOGGIN** | 0.81 | 0.55 | |

**p-values of correlation matrix**

| | BMP2 | BMP4 | NOGGIN |
|---|---|---|---|
| **BMP2** | | <0.001 | <0.001 |
| **BMP4** | <0.001 | | <0.001 |
| **NOGGIN** | <0.001 | <0.001 | |

**Figure S6:** Expression profiles for BMP2/4 and NOGGIN mRNA in DLGG databases. (A) RNA levels for BMP2, BMP4 and NOGGIN in non-tumoral brain, oligodendrogliomas, astrocytomas and oligoastrocytomas in Rembrandt [3] and Kamoun [4] databases. Significance was tested using multiple t-tests (p-values were corrected with Bonferroni correction for multiple comparisons). The number of cases is indicated in brackets. (B) Pearson correlation matrix of mRNA expression of BMP2, BMP4 and NOGGIN in the low grade glioma TCGA database reveals significant correlation. These histograms, scatter dot diagrams and correlation matrix were created with the glioma database mining website GLIOVIS [5].

QPCR analysis of BMP2, BMP4 and NOGGIN expression in DLGG

**A**

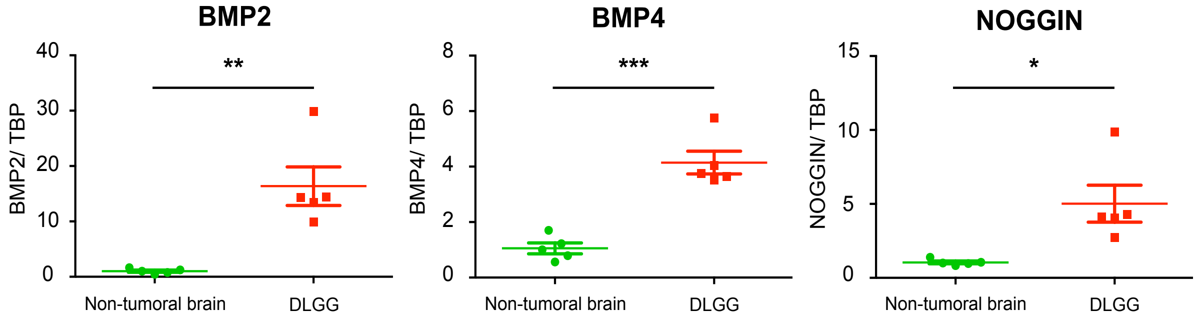


**B**

**Correlation matrix (Pearson, n=5 cases)**

| | BMP2 | BMP4 | NOGGIN |
|---|---|---|---|
| **BMP2** | | 0.91 | 0.95 |
| **BMP4** | 0.91 | | 0.98 |
| **NOGGIN** | 0.95 | 0.98 | |

**p values of correlation matrix**

| | BMP2 | BMP4 | NOGGIN |
|---|---|---|---|
| **BMP2** | | 0.031 | 0.015 |
| **BMP4** | 0.031 | | 0.003 |
| **NOGGIN** | 0.015 | 0.003 | |

**C**

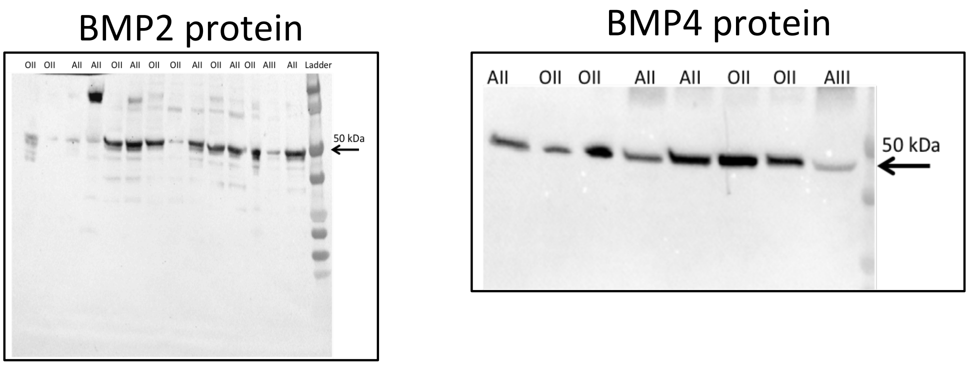


**Figure S7:** Expression of BMP2/4 and NOGGIN in DLGG. (A) QPCR data showing increased expression of BMP2, BMP4 and NOGGIN in five IDH1 R132H mutated DLGG resections (4 grade II oligodendrogliomas and 1 astrocytoma) compared to non-tumoral human brain tissues (epileptic tissues). Tests are two-tailed t-tests (n=5). (B) Pearson correlation matrix reveals that BMP2, BMP4 and NOGGIN expression are significantly correlated in these DLGG samples. (C) Western blot analysis for BMP2 and BMP4 proteins extracted from IDH1-mutated grade II oligodendrogliomas (OII) and IDH1-mutated grade III or II astrocytomas (AIII, AII). The same amount of proteins (20 µg) per lane was loaded by lane.

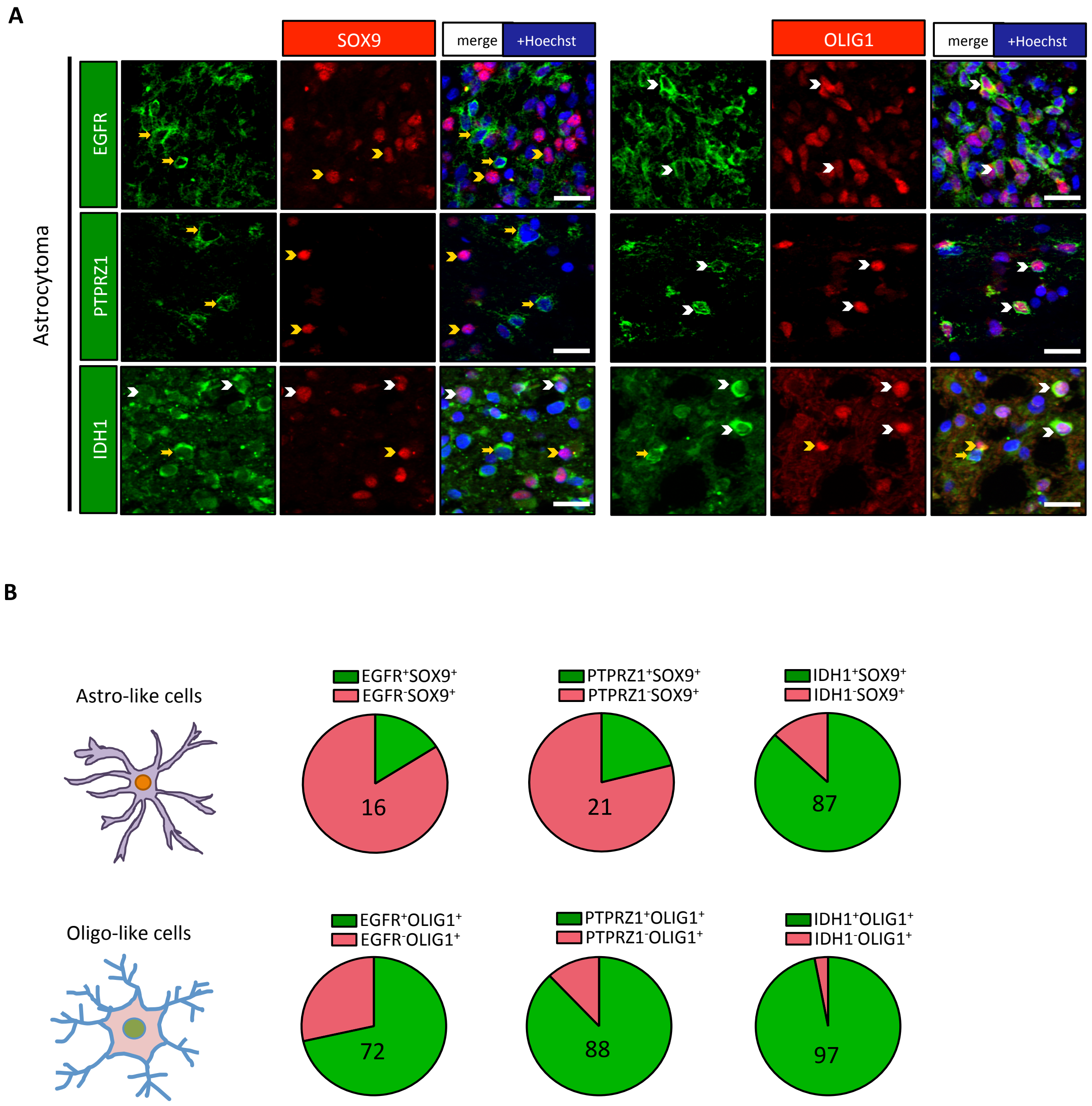


**Figure S8:** $OLIG1^{+}$ cells specifically express/activate various receptors and IDH1. Double immunofluorescences for indicated proteins on one astrocytoma. White arrowheads show double positive cells while yellow arrowheads/arrows show single positive cells. Scale bars=20 µm. (A) $OLIG1^{+}$ cells preferentially express EGFR and PTPRZ1 compared to $SOX9^{+}$ cells. Expression of IDH1 was closely associated to $OLIG1^{+}$ cells. (B) Pie diagrams represent the percentage of double positive (green) and single positive (red) cells in astro-like $SOX9^{+}$ (upper lane) and oligo-like $OLIG1^{+}$ (lower lane) cell populations. Numbers indicate the percentage of double positive cells.

Single cell RNA seq analysis of IDH1 gene expression in adult mouse and human brain

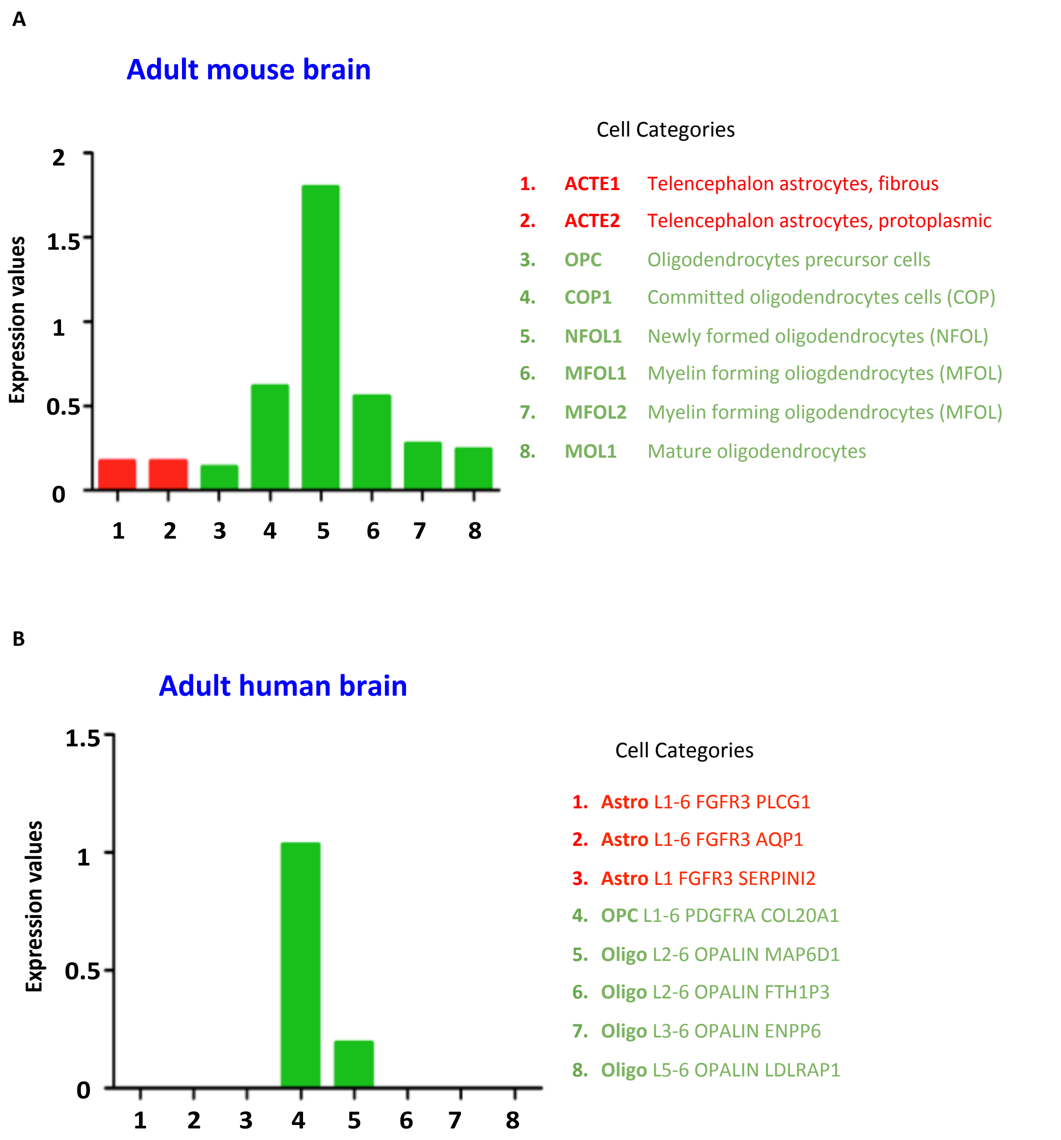


**Figure S9:** Cell type specific expression of IDH1 in adult mouse and human brain. Single cell transcriptomic analysis performed in the adult mouse and human brain reveals oligodendrocytic specific expression of IDH1 gene. (A) In mice, oligodendrocytic cells at different differentiation stages express IDH1 with highest expression in newly formed oligodendrocytes while expression in astrocytes is very low. (B) In the human brain, highest expression is observed in PDGFRA$^+$ oligodendrocyte cells (OPC). Mouse and human brain single cell transcriptomics data used to draw these histograms were obtained from the mousebrain.org [6] and Allen brain (celltypes.brain-map.org/rnaseq/human_m1_10x) [2] databases respectively. The different cell categories are those specified in the database websites.

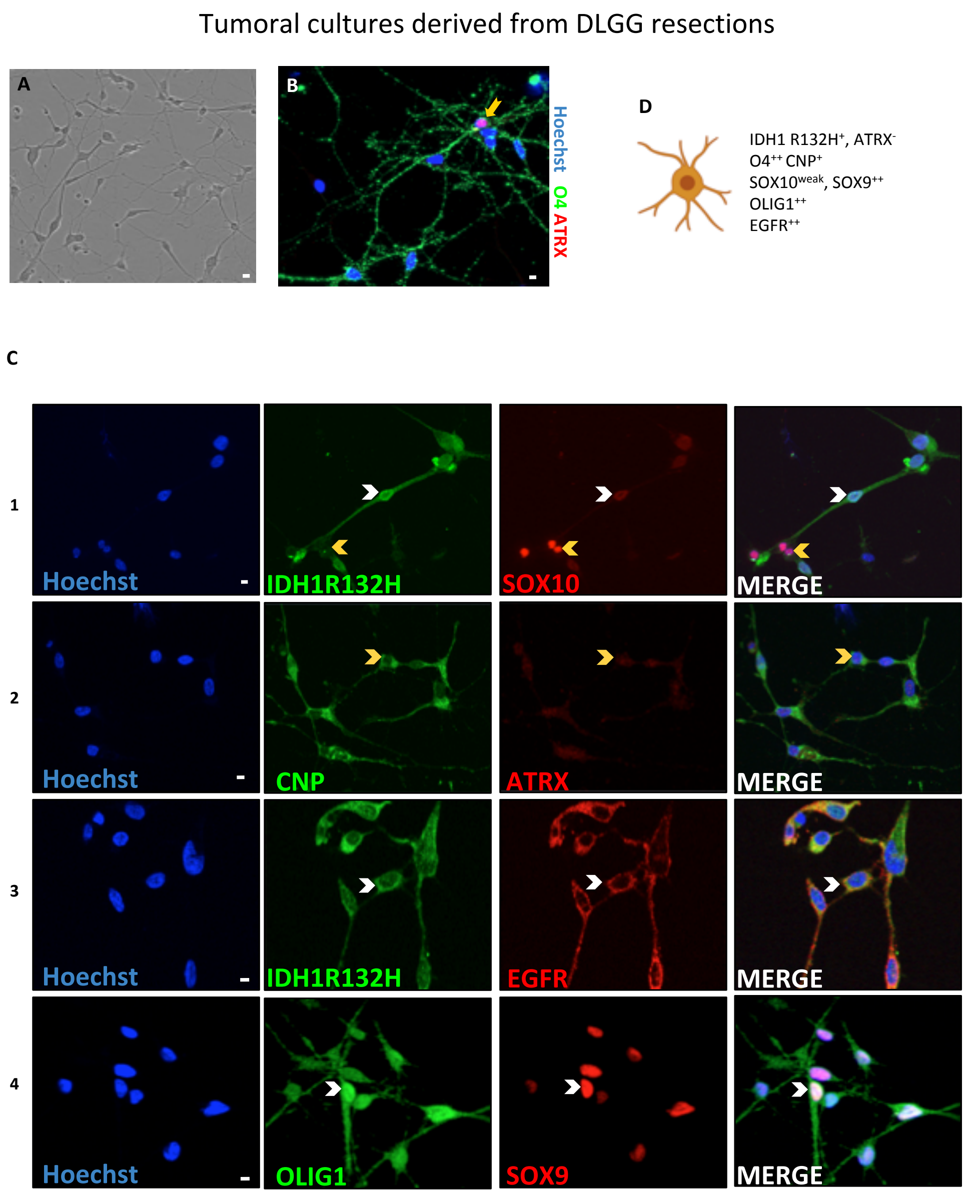


**Figure S10:** Phenotypic characterization of tumoral O4$^{+}$ cells isolated from DLGG resections. Double immunofluorescence for indicated proteins in a representative culture. (A) Morphology of O4-purified tumoral human cells in culture. Scale bar=10 µm. (B) Double stainings for O4 and ATRX show that the vast majority of cells (>90%) express O4 but not nuclear ATRX and are thus tumoral. One example of a rare O4$^{+}$ cell maintaining expression of ATRX, and thus probably non-tumoral, is presented to show the quality of staining (yellow arrow). Scale bar=5 µm. (C) Positive staining for IDH1 R132H (lane 1) and absence of nuclear ATRX (lane 2) confirmed the tumoral status of the culture. IDH1 R132H$^{+}$ cells express SOX10 at low level (white arrowhead, lane 1) in contrast to the rare IDH1 R132H- non-tumoral cells, which were highly positive for SOX10 (yellow arrowhead, lane 1). Tumoral cells also express EGFR (lane 3), the oligodendrocytic markers CNP (lane 2) and OLIG1 together with SOX9 (white arrowheads) (lane 4). Note that OLIG1 is localized both in the nucleus and cytoplasm. Scale bars=5 µm (D) Graphical summary of the phenotype of tumoral O4-purified cell.

Non-tumoral cultures derived from DLGG resections

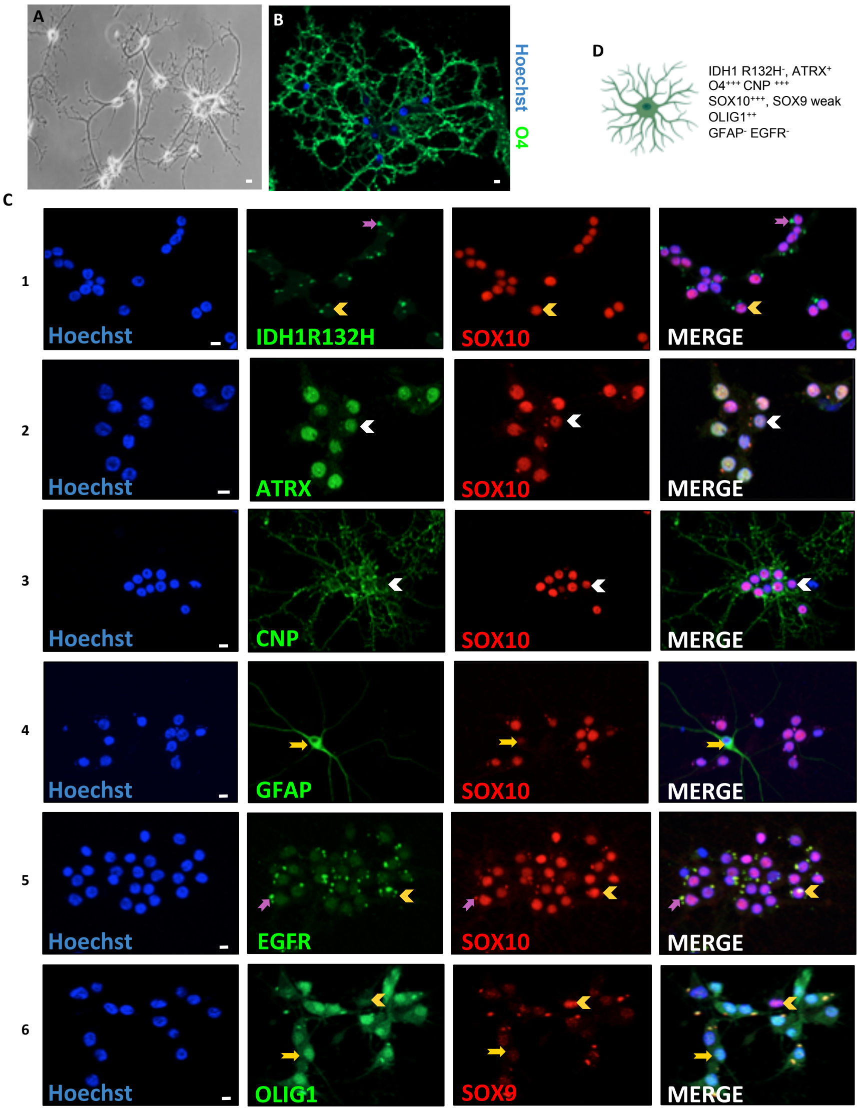


**Figure S11:** Phenotypic characterization of non-tumoral O4$^{+}$ cells isolated from DLGG samples. (A) Morphology of highly branched non-tumoral cells isolated with O4 purification from a supratotal glioma resection with ATRX and IDH1 R132H mutations. Scale bar=10 µm. (B) Immunofluorescence for O4 shows the very high expression of this antigen. The vast majority of cells (>90%) are O4$^{+}$. (C) Representative images of immunofluorescences performed on O4-purified cells. White arrowheads represent double positive cells while yellow arrowheads/arrows indicate single positive cells. Of note, non-tumoral cells isolated from DLGG resections often-present highly fluorescent green and red auto fluorescent cytoplasmic aggregates (most likely lipofuscin) (pink arrows), which we considered as non-specific signals. Lane 1 and 2: absence of IDH1 R132H protein expression and strong nuclear staining for ATRX show the non-tumoral status of these cells. Lane 3, 4 and 5: these cells are highly positive for oligodendrocytic markers such as CNP, OLIG1, and SOX10 and have small and perfectly round nuclei. Lane 4, 5 and 6: in contrast, the vast majority of these cells (>90%) do not express EGFR, GFAP and SOX9. Two examples of very rare GFAP$^{+}$ SOX10$^{-}$ (lane 4) and SOX9$^{+}$ OLIG1$^{-}$ (lane 6) cells are presented to show the quality of immunofluorescence (yellow arrow). Scale bars=10 µm (D) Graphical summary of the phenotype of non-tumoral human O4-purified cells.

LGG 275 cell line

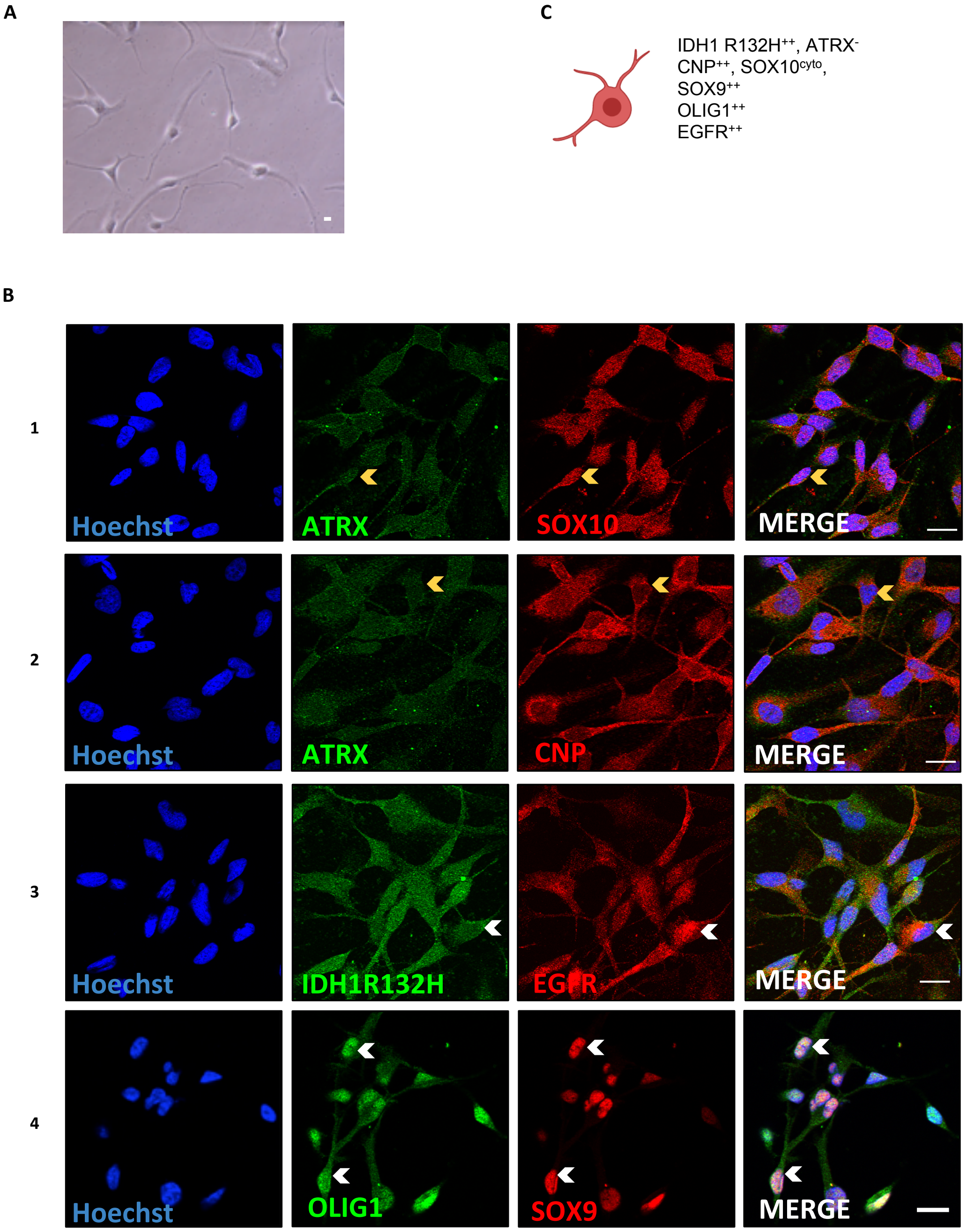


**Figure S12:** Phenotypic characterization of the LGG275 cell line. (A) Morphology of LGG275 cells. Scale bar=10 µm. (B) Double immunofluorescences for indicated proteins. The tumoral status of these cells is indicated by the detection of IDH1 R132H and absence of nuclear ATRX (lane 1, 2, 3). LGG275 cells show staining for CNP, EGFR, OLIG1 and SOX9 (lane 2, 3, 4). Detection of SOX10 is weak in these cells (lane 1). Note that LGG275 cells show different nuclei size and shape. White arrowheads represent double positive cells while yellow arrowheads indicate single positive cells. Scale bars=20 µm

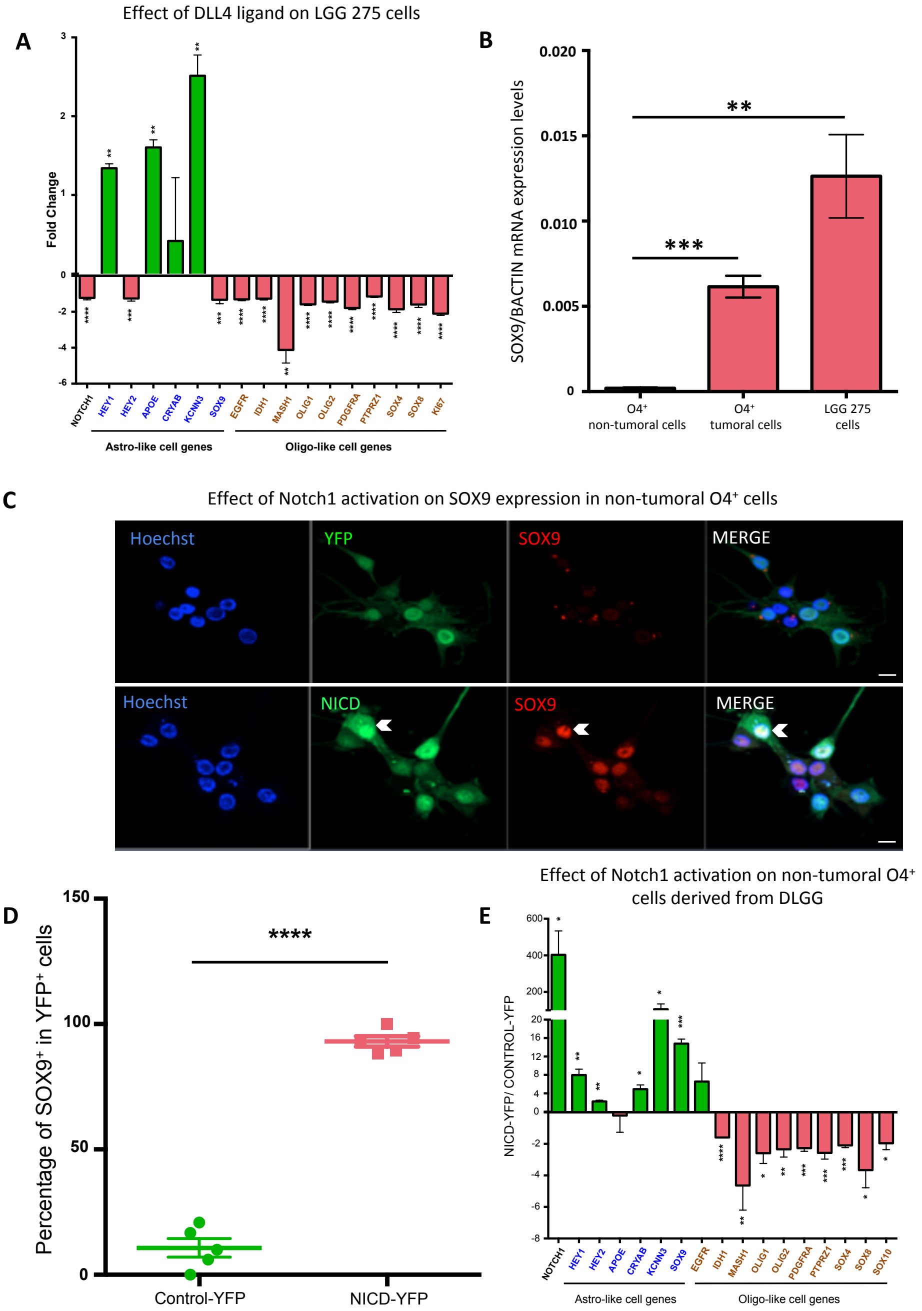


**Figure S13:** DLL4 effect on cell phenotype and regulation of SOX9 by Notch1 signalling. (A) QPCR analysis for 17 genes on LGG275 cells exposed to DLL4 ligand for 5 days. Test=two-tailed t-test, n=3 independent experiments. (B) SOX9 mRNA levels measured by QPCR in O4-purified non-tumoral cells, O4-purified tumoral cells and LGG275 cells. Test=two-tailed t-test, n=3 independent experiments. (C) Immunofluorescence for SOX9 in O4-purified non-tumoral cells transduced with YFP and YFP-NICD lentiviruses. Scale bars=10 µm. (D) Quantification of immunofluorescence presented in (C), n=6 fields, 3 coverslips. Test=two-tailed t-test. (E) QPCR analysis for indicated genes in non-tumoral cells purified with O4 from DLGG resections (n=3 patients). Values represent the mean $^{+}$/-S.E.M of gene expression fold change observed in cells transduced with NICD-YFP vs. YFP lentiviruses. Genes in blue and brown are markers found preferentially associated with respectively SOX9$^{+}$ and OLIG1$^{+}$ cells on DLGG sections. Tests=two-tailed t-tests.

**A**

CRYAB, HEY2 and BMP2 co-expression in TCGA_LGG dataset

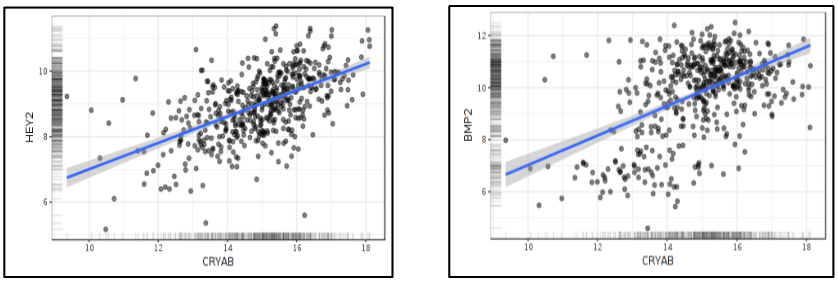


**Correlation matrix (Pearson, n=511 cases)**

| | CRYAB | HEY2 | BMP2 |
|---|---|---|---|
| **CRYAB** | | 0.56 | 0.50 |
| **HEY2** | 0.56 | | 0.27 |
| **BMP2** | 0.50 | 0.27 | |

**p values of correlation matrix**

| | CRYAB | HEY2 | BMP2 |
|---|---|---|---|
| **CRYAB** | | <0.001 | <0.001 |
| **HEY2** | <0.001 | | <0.001 |
| **BMP2** | <0.001 | <0.001 | |

**B**

HEY1 in TCGA_GBMLGG dataset

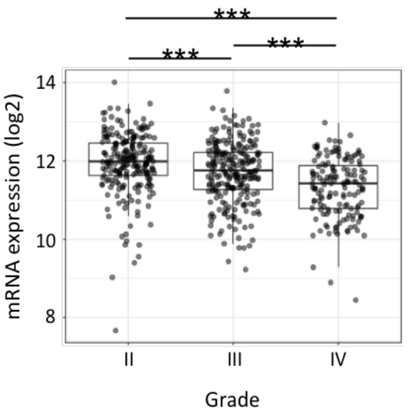


HEY2 in TCGA_GBM-LGG dataset

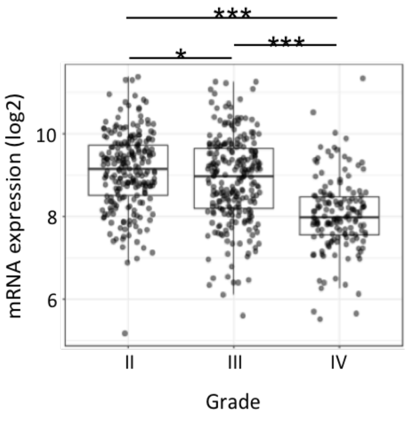


**Figure S14:** Database mining for HEY1/2 expression and correlation in different glioma grades. (A) Upper panels: Expression of CRYAB is highly correlated to HEY2 and BMP2 expression in the diffuse low-grade gliomas TCGA database. Lower panels: Pearson correlation matrix and p-values for CRYAB, HEY2 and BMP2. These scatter dot diagrams and correlation matrix were created with the glioma database mining website GLIOVIS [5]. (B) Expression of HEY1 and HEY2 expression is reduced in high grade compared to low-grade gliomas in the TCGA database. Significance was tested using multiple t-tests (p-values were corrected with Bonferroni correction for multiple comparisons). Box plots were obtained from the glioma database mining website GLIOVIS [5].

**Table S1:** Detailed information of the patients used in the article

| **Tumor** | **Techniques** | **Figure** | **Age** | **Gender** | **Diagnosis (subtype, grade)** | **IDH1 mutation** | **ATRX** | **1p/19q deletion** |
|---|---|---|---|---|---|---|---|---|
| Patient 1 LGG 236 | IF | 1, S2B, 2, 3, 4, | 49 | F | Oligodendroglioma II | IDH1R132H | Preserved | Co-deleted |
| Patient 2 LGG 253 | IF | S2A | 26 | M | Oligodendroglioma II | IDH1R132H | Preserved | Co-deleted |
| Patient 3 LGG 39 | IF | S2A | 67 | M | Oligodendroglioma II | IDH1R132H | Preserved | Co-deleted |
| Patient 4 LGG 234 | IF | 1, S2B, S3, S4, S5, S8 | 32 | M | Astrocytoma Grade II | IDH1R132H | Loss | No co-deletion |
| Patient 5 LGG 244 | IF | S2A | 40 | M | Astrocytoma II | IDH1R132H | Loss | No co-deletion |
| Patient 6 LGG 188 | IF | S2A | 42 | F | Astrocytoma II | IDH1R132H | Loss | No co-deletion |
| LGG 270 | Tumoral cells qPCR-YFP, NICD | 5A, S10 | 38 | M | Astrocytoma II | IDH1R132H | Loss | No co-deletion |
| LGG275 | Cell line | 5B-E, 6A-E, S12, S13A | 40 | F | Astrocytoma II | IDH1R132H | Loss | No co-deletion |
| LGG 289 | Tumoral cells qPCR-YFP, NICD, Cell characterization | 5A, S10 | 47 | M | Astrocytoma II | IDH1R132H | Loss | No co-deletion |
| LGG 312 | Tumoral cells qPCR-YFP, NICD | 5A, S10 | 31 | F | Astrocytoma II | IDH1R132H | Loss | No co-deletion |
| LGG 318 | Tumoral cells qPCR-YFP, NICD | 5A, S10 | 26 | M | Astrocytoma II | IDH1R132G | Loss | No co-deletion |
| LGG 307 | Non-tumoral | S11, | 52 | F | Oligodendroglioma II | IDH1wt | No Loss | Co-deleted |

| | | | | | | | | |
|---|---|---|---|---|---|---|---|---|
| | cells qPCR-YFP, NICD | S13B-E | | | | | | |
| LGG 335 | Non-tumoral cells qPCR-YFP, NICD<br>Cell characterization | S11, S13B-E | 30 | M | Oligodendroglioma II | IDH1R132H | No Loss | Co-deleted |
| LGG 338 | Non-tumoral cells qPCR-YFP, NICD<br>Cell characterization | S11, S13B-E | 41 | F | Astrocytoma II | IDH1R132H | Loss | No co-deletion |

**Table S2:** List of cellular markers and supporting references for their specific expression.

| Protein Name (short form) | Full form | Astrocyte/Oligodendrocyte/progenitor cell | Functions | Supporting references |
|---|---|---|---|---|
| APOE | Apolipoprotein E | Astrocytes | | [7] |
| CRYAB | Alpha-crystallin B chain | | | [8,9] |
| GPR17 | G Protein-Coupled Receptor 17 | Oligodendrocytes | GPR17 is a downstream target of OLIG2, regulates oligodendrocyte survival and is involved in timing myelination | [10,11] |
| ID4 | Inhibitor of differentiation 4 | | ID4 is expressed in cells with astrocytic features in oligodendroglioma and astrocytomas | [12,13] |
| KCNN3/SK3/KCa2.3 | Small conductance calcium-activated potassium channel 3 | | KCNN3 is upregulated in diffuse astrocytoma<br><br>KCNN3 is expressed by GFAP positive astrocytes | [14]<br><br>[15] |
| MASH1/ ASCL1 | Achaete-scute homolog 1 | | | [16-18] |
| OLIG1 | Oligodendrocyte Transcription Factor 1 | Oligodendrocytes | | [19] |

| | | | | |
|---|---|---|---|---|
| PDGFRA | Platelet-derived growth factor receptor alpha | | OPC marker | [20,21] |
| PTPRZ1 | Receptor-type tyrosine-protein phosphatase zeta | Oligodendrocytes | | [22] |
| SOX8 | SRY-Box Transcription Factor 8 | Oligodendrocytes | Specific marker of oligodendrocytes according to Ben Barres database | [23]<br><br>[1] |
| SOX9 | SRY-Box Transcription Factor 9 | Astrocytes | | [24] |

**Table S3**: PCR primer pairs used for quantitative RT-PCR.

| Gene | Forward primer | Reverse primer |
|---|---|---|
| ACTB | GGACTTCGAGCAAGAGATGG | AGCACTGTGTTGGCGTACAG |
| APOE | GGTCGCTTTTGGGATTACCT | TTCCTCCAGTTCCGATTTGT |
| BMP2 | TGTATCGCAGGCACTCAGGTCA | CCACTCGTTTCTGGTAGTTCTTC |
| BMP4 | CTGGTCTTGAGTATCCTGAGCG | TCACCTCGTTCTCAGGGATGCT |
| CRYAB | TCATCTCCAGGGAGTTCCAC | AGGACCCCATCAGATGACAG |
| EGFR | GTGTGCCCACTACATTGACG | CTTCCAGACCAGGGTGTTGT |
| GPR17 | GGAAGAACAACCCCTGAACA | TCCCTCTTCTGGGTCATTTG |
| HEY1 | TGTTTGGTTCAAGGCAGCTC | TGATGCACTGCTGGATGGTA |
| HEY2 | GCACCCTGAAGGTAGCCATA | AGTTACCGAGCTGCCTTGAA |
| IDH1 | AGTCTGCAAGACTGGGAGGA | CAGAACCGCCACTGATTTTT |
| KCNN3 | CTGCCGCCAAAATAAACATT | GCCTGGCACAAGCTTTCTAC |
| MASH1 | CAAGAGAGCGCAGCCTTAGT | CTGGCGCCTTCTTGTTTCTA |
| MKI67 | CCCCCACCAGAACTAACAGA | ACTTTGATGCCCTCATCACC |
| NOGGIN | TCGAACACCCAGACCCTATC | ATGAAGCCTGGGTCGTAGTG |
| NOTCH1 | TCCACCAGTTTGAATGGTCA | CGCAGAGGGTTGTATTGGTT |
| OLIG1 | CGCAGAGAGTTTTCGCTCTT | GCGGTTGGTTTTCGTTTTTA |
| OLIG2 | GACAAGCTAGGAGGCAGTGG | CGGCTCTGTCATTTGCTTCT |
| PDGFRA | GCTGATCCGTGCTAAGGAAG | CGACCAAGTCCAGAATGGAT |
| PTPRZ1 | CAATCGCATAGGGACGAAAT | AGTGACTGGTTGGGAAGTGG |
| SOX10 | AGCCCAGGTGAAGACAGAGA | TGTAGGCGATCTGTGAGGTG |
| SOX4 | GCACTAGGACGTCTGCCTTT | ACACGGCATATTGCACAGGA |
| SOX8 | TGATTCACCTGCACTGCTTC | AGCAACTTCTCGGCTGTGTT |
| SOX9 | GGAATGTTTCAGCAGCCAAT | TGGTGTTCTGAGAGGCACAG |

**Table S4**: Antibodies used for immunostaining.

| Antibody | Species | References | Suppliers | Dilution |
|---|---|---|---|---|
| Alpha B Crystallin (CRYAB) | Rabbit | 15808-1-AP | Proteintech | 1:300 |
| APOE | Mouse | sc-13521 | Santa Cruz | 1:200 |
| ATRX | Rabbit | HPA-001906 | Sigma | 1:500 |
| BMP4 | Rabbit | NBP1-95882 (EPR6211) | Novus Biological | 1:100 |
| CNPase | Mouse | C5922 | Sigma | 1:100 |
| EGFR | Rabbit | 4267 | Cell Signaling | 1:50 |
| GFAP | Rabbit | Z0334 | Dako | 1:5000 |
| GPR17 | Rabbit | Gift from Davide Lecca | | 1:100 |
| HEY1 | Rabbit | GTX118007 | GeneTex | 1:200 |
| HEY2 | Rabbit | 10597-I-AP | Proteintech | 1:300 |
| ID4 | Rabbit | BCH-9/82-12 | Biocheck | 1:200 |
| IDH1 | Rabbit | 8137 | Cell Signaling | 1:500 |
| IDH1R132H | Mouse | DIA-H09 | Dianova | 1:100 |
| KCNN3 (SK3) | Rabbit | APC-025 | Alomone Labs | 1:1000 |
| MASH1 | Mouse | 556604 | BD Biosciences | 1:100 |
| MASH1 | Rabbit | ab211327 | Abcam | 1:100 |
| MKI67 | Mouse | 556003 | BD Pharmingen | 1:500 |
| NICD | Sheep | AF3647 | R&D Systems | 1:200 |
| O4 | Mouse | | supernatant from hybridoma | 1:2 |
| OLIG1 | Goat | AF2417 | R&D Systems | 1:300 |
| p-ERK | Rabbit | 9910 | Cell Signaling | 1:200 |

| | | | | |
|---|---|---|---|---|
| p-SMAD1/5 | Rabbit | 9516 | Cell Signaling | 1:800 |
| PDGFRa | Rabbit | 3174 | Cell Signaling | 1:200 |
| PTPRz1 | Rabbit | sc-25432 | Santa Cruz | 1:200 |
| SOX10 | Mouse | MAB2864 | R&D Systems | 1:200 |
| SOX10 | Rabbit | ab155279 | Abcam | 1:200 |
| SOX2 | Rabbit | 23064 | Cell Signaling | 1:200 |
| SOX4 | Mouse | AMab91378 | Atlas | 1:250 |
| SOX8 | Rabbit | 20627-1-AP | Proteintech | 1:300 |
| SOX9 | Rabbit | 82630S | Cell Signaling | 1:300 |
| SOX9 | Goat | AF3075-SP | R&D Systems | 1:300 |

### References for supplementary information

1. Zhang, Y.; Sloan, S.A.; Clarke, L.E.; Caneda, C.; Plaza, C.A.; Blumenthal, P.D.; Vogel, H.; Steinberg, G.K.; Edwards, M.S.; Li, G., et al. Purification and Characterization of Progenitor and Mature Human Astrocytes Reveals Transcriptional and Functional Differences with Mouse. *Neuron* **2016**, *89*, 37-53, doi:10.1016/j.neuron.2015.11.013.
2. Hodge, R.D.; Bakken, T.E.; Miller, J.A.; Smith, K.A.; Barkan, E.R.; Graybuck, L.T.; Close, J.L.; Long, B.; Johansen, N.; Penn, O., et al. Conserved cell types with divergent features in human versus mouse cortex. *Nature* **2019**, *573*, 61-68, doi:10.1038/s41586-019-1506-7.
3. Madhavan, S.; Zenklusen, J.C.; Kotliarov, Y.; Sahni, H.; Fine, H.A.; Buetow, K. Rembrandt: helping personalized medicine become a reality through integrative translational research. *Mol Cancer Res* **2009**, *7*, 157-167, doi:10.1158/1541-7786.MCR-08-0435.
4. Kamoun, A.; Idbaih, A.; Dehais, C.; Elarouci, N.; Carpentier, C.; Letouze, E.; Colin, C.; Mokhtari, K.; Jouvet, A.; Uro-Coste, E., et al. Integrated multi-omics analysis of oligodendroglial tumours identifies three subgroups of 1p/19q co-deleted gliomas. *Nat Commun* **2016**, *7*, 11263, doi:10.1038/ncomms11263.
5. Bowman, R.L.; Wang, Q.; Carro, A.; Verhaak, R.G.; Squatrito, M. GlioVis data portal for visualization and analysis of brain tumor expression datasets. *Neuro Oncol* **2017**, *19*, 139-141, doi:10.1093/neuonc/now247.
6. La Manno, G.; Siletti, K.; Furlan, A.; Gyllborg, D.; Vinsland, E.; Langseth, C.M.; Khven, I.; Johnsson, A.; Nilsson, M.; Lönnerberg, P., et al. Molecular architecture of the developing mouse brain. *bioRxiv* **2020**, 10.1101/2020.07.02.184051, 2020.2007.2002.184051, doi:10.1101/2020.07.02.184051.
7. Pitas, R.E.; Boyles, J.K.; Lee, S.H.; Foss, D.; Mahley, R.W. Astrocytes synthesize apolipoprotein E and metabolize apolipoprotein E-containing lipoproteins. *Biochim Biophys Acta* **1987**, *917*, 148-161, doi:10.1016/0005-2760(87)90295-5.
8. Avliyakulov, N.K.; Rajavel, K.S.; Le, K.M.; Guo, L.; Mirsadraei, L.; Yong, W.H.; Liau, L.M.; Li, S.; Lai, A.; Nghiemphu, P.L., et al. C-terminally truncated form of alphaB-crystallin is associated with IDH1 R132H mutation in anaplastic astrocytoma. *J Neurooncol* **2014**, *117*, 53-65, doi:10.1007/s11060-014-1371-z.
9. Goplen, D.; Bougnaud, S.; Rajcevic, U.; Boe, S.O.; Skaftnesmo, K.O.; Voges, J.; Enger, P.O.; Wang, J.; Tysnes, B.B.; Laerum, O.D., et al. alphaB-crystallin is elevated in highly infiltrative apoptosis-resistant glioblastoma cells. *Am J Pathol* **2010**, *177*, 1618-1628, doi:10.2353/ajpath.2010.090063.
10. Chen, Y.; Wu, H.; Wang, S.; Koito, H.; Li, J.; Ye, F.; Hoang, J.; Escobar, S.S.; Gow, A.; Arnett, H.A., et al. The oligodendrocyte-specific G protein-coupled receptor GPR17 is a cell-intrinsic timer of myelination. *Nat Neurosci* **2009**, *12*, 1398-1406, doi:10.1038/nn.2410.
11. Ou, Z.; Sun, Y.; Lin, L.; You, N.; Liu, X.; Li, H.; Ma, Y.; Cao, L.; Han, Y.; Liu, M., et al. Olig2-Targeted G-Protein-Coupled Receptor Gpr17 Regulates Oligodendrocyte Survival in Response to Lysolecithin-Induced Demyelination. *J Neurosci* **2016**, *36*, 10560-10573, doi:10.1523/JNEUROSCI.0898-16.2016.
12. Liang, Y.; Bollen, A.W.; Nicholas, M.K.; Gupta, N. Id4 and FABP7 are preferentially expressed in cells with astrocytic features in oligodendrogliomas and oligoastrocytomas. *BMC Clin Pathol* **2005**, *5*, 6, doi:10.1186/1472-6890-5-6.
13. Samanta, J.; Kessler, J.A. Interactions between ID and OLIG proteins mediate the inhibitory effects of BMP4 on oligodendroglial differentiation. *Development* **2004**, *131*, 4131-4142, doi:10.1242/dev.01273.
14. Rorive, S.; Maris, C.; Debeir, O.; Sandras, F.; Vidaud, M.; Bieche, I.; Salmon, I.; Decaestecker, C. Exploring the distinctive biological characteristics of pilocytic and low-grade diffuse astrocytomas using microarray gene expression profiles. *J Neuropathol Exp Neurol* **2006**, *65*, 794-807, doi:10.1097/01.jnen.0000228203.12292.a1.
15. Armstrong, W.E.; Rubrum, A.; Teruyama, R.; Bond, C.T.; Adelman, J.P. Immunocytochemical localization of small-conductance, calcium-dependent potassium channels in astrocytes of the rat supraoptic nucleus. *J Comp Neurol* **2005**, *491*, 175-185, doi:10.1002/cne.20679.
16. Nakatani, H.; Martin, E.; Hassani, H.; Clavairoly, A.; Maire, C.L.; Viadieu, A.; Kerninon, C.; Delmasure, A.; Frah, M.; Weber, M., et al. Ascl1/Mash1 promotes brain oligodendrogenesis during myelination and remyelination. *J Neurosci* **2013**, *33*, 9752-9768, doi:10.1523/JNEUROSCI.0805-13.2013.

17. Rousseau, A.; Nutt, C.L.; Betensky, R.A.; Iafrate, A.J.; Han, M.; Ligon, K.L.; Rowitch, D.H.; Louis, D.N. Expression of oligodendroglial and astrocytic lineage markers in diffuse gliomas: use of YKL-40, ApoE, ASCL1, and NKX2-2. *J Neuropathol Exp Neurol* **2006**, *65*, 1149-1156, doi:10.1097/01.jnen.0000248543.90304.2b.

18. Sugimori, M.; Nagao, M.; Parras, C.M.; Nakatani, H.; Lebel, M.; Guillemot, F.; Nakafuku, M. Ascl1 is required for oligodendrocyte development in the spinal cord. *Development* **2008**, *135*, 1271-1281, doi:10.1242/dev.015370.

19. Othman, A.; Frim, D.M.; Polak, P.; Vujicic, S.; Arnason, B.G.; Boullerne, A.I. Olig1 is expressed in human oligodendrocytes during maturation and regeneration. *Glia* **2011**, *59*, 914-926, doi:10.1002/glia.21163.

20. Ellison, J.A.; de Vellis, J. Platelet-derived growth factor receptor is expressed by cells in the early oligodendrocyte lineage. *J Neurosci Res* **1994**, *37*, 116-128, doi:10.1002/jnr.490370116.

21. Zhu, Q.; Zhao, X.; Zheng, K.; Li, H.; Huang, H.; Zhang, Z.; Mastracci, T.; Wegner, M.; Chen, Y.; Sussel, L., et al. Genetic evidence that Nkx2.2 and Pdgfra are major determinants of the timing of oligodendrocyte differentiation in the developing CNS. *Development* **2014**, *141*, 548-555, doi:10.1242/dev.095323.

22. Lamprianou, S.; Chatzopoulou, E.; Thomas, J.-L.; Bouyain, S.; Harroch, S. A complex between contactin-1 and the protein tyrosine phosphatase PTPRZ controls the development of oligodendrocyte precursor cells. *Proceedings of the National Academy of Sciences* **2011**, *108*, 17498-17503, doi:10.1073/pnas.1108774108.

23. Azar, S.; Leventoux, N.; Ripoll, C.; Rigau, V.; Goze, C.; Lorcy, F.; Bauchet, L.; Duffau, H.; Guichet, P.O.; Rothhut, B., et al. Cellular and molecular characterization of IDH1-mutated diffuse low grade gliomas reveals tumor heterogeneity and absence of EGFR/PDGFRalpha activation. *Glia* **2018**, *66*, 239-255, doi:10.1002/glia.23240.

24. Sun, W.; Cornwell, A.; Li, J.; Peng, S.; Osorio, M.J.; Aalling, N.; Wang, S.; Benraiss, A.; Lou, N.; Goldman, S.A., et al. SOX9 Is an Astrocyte-Specific Nuclear Marker in the Adult Brain Outside the Neurogenic Regions. *J Neurosci* **2017**, *37*, 4493-4507, doi:10.1523/JNEUROSCI.3199-16.2017.
